# Structural connectivity-based segmentation of the human entorhinal cortex

**DOI:** 10.1101/2021.07.16.452500

**Authors:** Ingrid Framås Syversen, Menno P. Witter, Asgeir Kobro-Flatmoen, Pål Erik Goa, Tobias Navarro Schröder, Christian F. Doeller

## Abstract

The medial (MEC) and lateral entorhinal cortex (LEC), widely studied in rodents, are well defined and characterized. In humans, however, the exact locations of their homologues remain uncertain. Previous functional magnetic resonance imaging (fMRI) studies have subdivided the human EC into posterior-medial (pmEC) and anterior-lateral (alEC) parts, but uncertainty remains about the choice of imaging modality and seed regions, in particular in light of a substantial revision of the classical model of EC connectivity based on novel insights from rodent anatomy. Here, we used structural, not functional imaging, namely diffusion tensor imaging (DTI) and probabilistic tractography to segment the human EC based on differential connectivity to other brain regions known to project selectively to MEC or LEC. We defined MEC as more strongly connected to presubiculum and retrosplenial cortex (RSC), and LEC as more strongly connected to distal CA1 and proximal subiculum (dCA1pSub) and orbitofrontal cortex (OFC). Although our DTI segmentation had a larger medial-lateral component than in previous fMRI studies, our results show that the human MEC and LEC homologues have a border oriented both towards the posterior-anterior and medial-lateral axes, supporting the differentiation between pmEC and alEC.

## Introduction

The entorhinal cortex (EC) is a part of the medial temporal lobe, and a central structure for memory formation and navigation (Eichenbaum, Yonelinas, & Ranganath, 2007; Moser & Moser, 2013; Suzuki & Eichenbaum, 2000). It is classically viewed as a hub for processing and relaying information from the neocortex to the hippocampus, and vice versa (Buzsáki, 1996; Lavenex & Amaral, 2000). The EC can be divided into two main subregions – ‘medial’ entorhinal cortex (MEC) and ‘lateral’ entorhinal cortex (LEC) – which differ in both functional properties and connectivity with other regions (Canto, Wouterlood, & Witter, 2008; Kerr, Agster, Furtak, & Burwell, 2007; van Strien, Cappaert, & Witter, 2009). Both the function and anatomy of the EC subregions have been widely studied in rodents and non-human primates. Based mainly on research in rodents, the MEC is associated with spatial processing in a global, allocentric frame of reference, given the prevalence of spatially modulated cells such as grid and head direction cells (Fyhn, Molden, Witter, Moser, & Moser, 2004; Hafting, Fyhn, Molden, Moser, & Moser, 2005; Høydal, Skytøen, Andersson, Moser, & Moser, 2019; Knierim, Neunuebel, & Deshmukh, 2014). In contrast, the LEC contains cells that are sensitive to the presence of objects in a local frame of reference or processing of time (Deshmukh & Knierim, 2011; Knierim et al., 2014; Tsao, Moser, & Moser, 2013; Tsao et al., 2018). However, although recent years have seen a stark increase in functional imaging studies of the human EC (Bellmund, Deuker, & Doeller, 2019; Chen, Vieweg, & Wolbers, 2019; Montchal, Reagh, & Yassa, 2019; Maass, Berron, Libby, Ranganath, & Düzel, 2015; Navarro Schröder, Haak, Zaragoza Jimenez, Beckmann, & Doeller, 2015; Reagh & Yassa, 2014; Schultz, Sommer, & Peters, 2012), the exact locations of the human homologues of MEC and LEC remain somewhat uncertain. This is an ongoing challenge for functional studies of the EC in humans and also makes it difficult to conduct translational research on the origins of neurodegenerative processes such as occurring in Alzheimer’s disease, which start in the EC and trans-entorhinal area (Braak & Braak, 1992).

In anatomical and functional studies of the human brain, magnetic resonance imaging (MRI) has become an invaluable tool. Functional MRI (fMRI) studies have shown that the properties of the rodent and non-human primate EC also apply to the human EC (Doeller, Barry, & Burgess, 2010; Reagh & Yassa, 2014; Schultz et al., 2012). Based on the subdivision of the rodent EC into MEC and LEC, studies have tried to localize their respective homologue regions in humans. Previous fMRI studies tested connectivity ‘fingerprints’ of EC subregions to other parts of the brain. Studies in rodents and non-human primates have demonstrated a largely similar organization of EC connectivity across species (Canto et al., 2008), thus predicting distinct fMRI connectivity fingerprints for the two subregions in humans as well. The resulting delineations of putative human homologue regions of the rodent MEC and LEC were labeled posteromedial EC (pmEC) and anterolateral EC (alEC), based on the outcome of two independent fMRI studies that tested local and global connectivity, respectively (Maass et al., 2015; Navarro Schröder et al., 2015). However, it remains unclear whether the results were affected by the nature of the imaging modality (fMRI) or the choice of seed brain regions used to identify the subregions.

In addition to the neuroimaging modality, the second reason for a re-evaluation has gained additional importance since the assumption about EC connectivity on which parts of the previous fMRI studies (Maass et al., 2015; Navarro Schröder et al., 2015) were based has been recently revised. For years, the existence of two parallel cortical connectivity streams through the EC has been the accepted model (Nilssen, Doan, Nigro, Ohara, & Witter, 2019; Ranganath & Ritchey, 2012; Witter, Doan, Jacobsen, Nilssen, & Ohara, 2017). This comprises one pathway into the hippocampus via the parahippocampal/postrhinal cortex (PHC/POR) and MEC (the “where” pathway), and a parallel pathway via the perirhinal cortex (PRC) and LEC (the “what” pathway). However, recent evidence substantially challenged this view. Doan and colleagues found that POR in rats, which corresponds to the PHC in humans, does also project to LEC. These authors further argue that existing data in monkeys substantiate this notion (Doan, Lagartos-Donate, Nilssen, Ohara, & Witter, 2019). This is in line with new findings in humans indicating that the hippocampal-entorhinal-neocortical connections are far more complex than a pure segregation into “where” and “what” pathways (C.-C. Huang, Rolls, Hsu, Feng, & Lin, 2021).

In order to more accurately identify the human homologues of MEC and LEC, we should take advantage of known unique connections to each subregion. For example, in rodents the presubiculum projects almost exclusively to MEC, whereas distal CA1 and proximal subiculum (dCA1pSub, i.e. the border region between CA1 and subiculum) project most strongly to LEC (Caballero-Bleda & Witter, 1993; Honda & Ishizuka, 2004; Witter & Amaral, 1991; Witter & Amaral, 2021). Meanwhile, the retrosplenial cortex (RSC) and the posterolateral orbitofrontal cortex (OFC) are respectively selectively connected with MEC or LEC (Hoover & Vertes, 2007; Jones & Witter, 2007; Kondo & Witter, 2014; Saleem, Kondo, & Price, 2008; Witter & Amaral, 2021; Wyss & Van Groen, 1992). To investigate the connectivity between these regions, there are several imaging modalities available. An alternative method to the widely used fMRI functional connectivity is to instead study structural connectivity using diffusion tensor imaging (DTI), another type of MRI (Powell et al., 2004; Zeineh, Holdsworth, Skare, Atlas, & Bammer, 2012). Here, one exploits the diffusion of water molecules inside white matter tracts and uses this to map the paths of these fibers – so-called tractography (Mori, Crain, Chacko, & van Zijl, 1999; Mori & Zhang, 2006). Mapping DTI connectivity from cortices that project selectively to either EC subregion could provide a novel line of evidence to identify MEC and LEC (Ezra, Faull, Jbabdi, & Pattinson, 2015; Máté et al., 2018; Saygin, Osher, Augustinack, Fischl, & Gabrieli, 2011).

The objective of this study is therefore to identify the human homologues of the rodent MEC and LEC using DTI, incorporating the novel insights from rodent anatomy. To achieve this, we performed probabilistic tractography on high-quality DTI data acquired by the Human Connectome Project (Fan et al., 2016). We identify the EC subregions by analyzing the connectivity from regions of interest (ROIs) that project selectively to either of them and compare these to the results from previous fMRI studies.

## Results

To visualize the connectivity paths between the EC and the regions hypothesized to be connected with its subregions, we ran probabilistic tractography between the regions. By seeding paths from all voxels in the EC, presubiculum, dCA1pSub, RSC and OFC ROIs, maps of the connectivity paths between the EC and the other ROIs were created. The resulting group averaged paths are shown in Figure 1. In all figures, blue color schemes are used for MEC-related regions, i.e. presubiculum and RSC, while red color schemes are used for LEC-related regions, i.e. dCA1pSub and OFC. The maps show that all the regions exhibit clear connectivity with the EC. Connections with dCA1pSub extend further anteriorly in the EC than the connections with the presubiculum, and the connections with presubiculum and RSC seem to take a similar route to the EC. The paths between OFC and EC, however, stand out from the others as they take a more lateral route, but the inferior part seems to pass close to dCA1pSub. Note that the colormap intensity in these maps does not represent the actual number of white matter tracts, but instead scales with the probability that the true path between the ROIs lies in that point. Corresponding connectivity paths for one example participant are shown in Figure 1–figure supplement 1.

**Figure 1:**
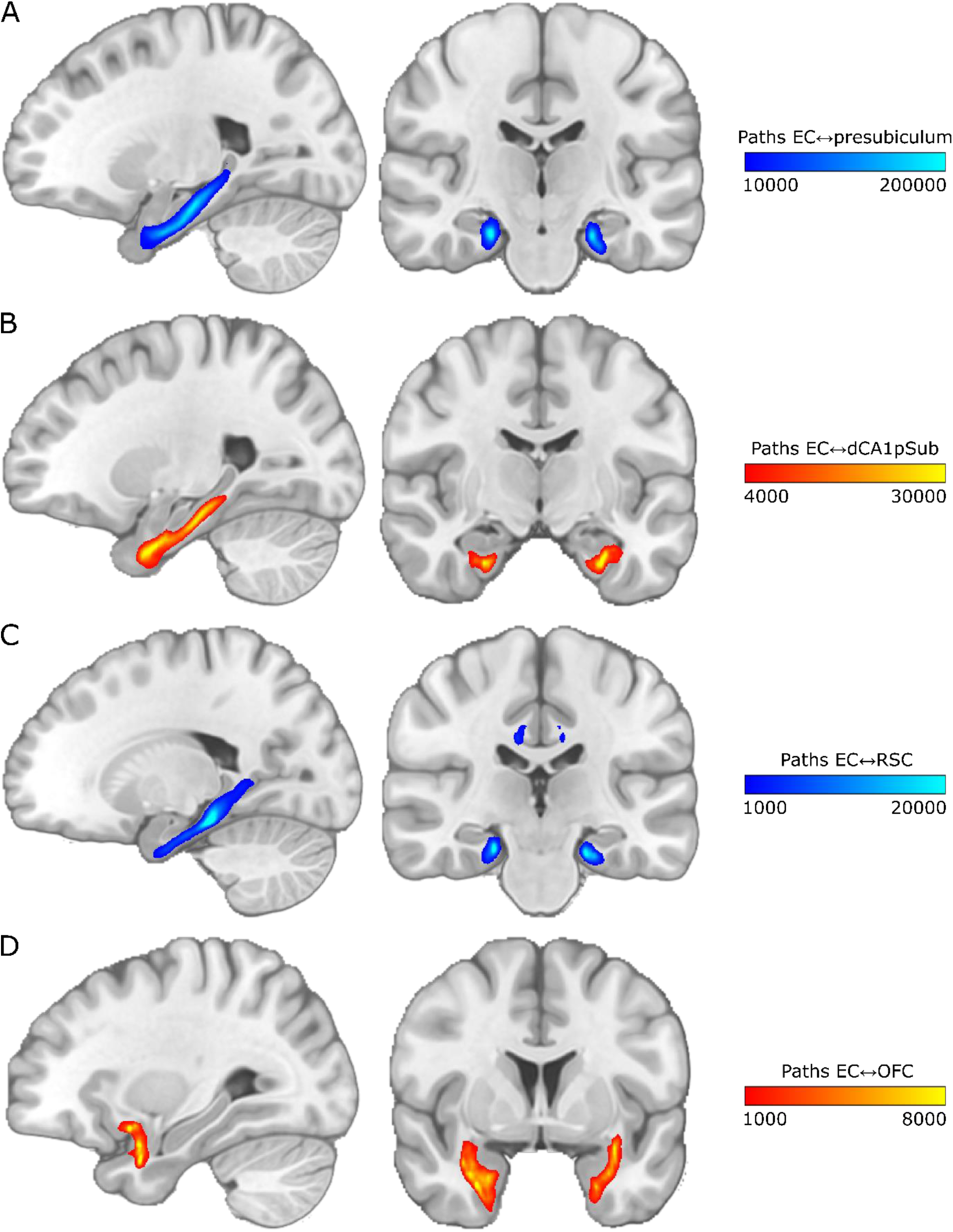
Group average connectivity paths between EC and presubiculum, dCA1pSub, RSC and OFC. Connectivity patterns are shown on sagittal (left) and coronal (right) slices in MNI space. The colormap intensity represents the number of probabilistic paths running through that voxel. **A:** Paths between EC and presubiculum, **B:** Paths between EC and dCA1pSub, **C:** Paths between EC and RSC, **D:** Paths between EC and OFC.

Because we wanted to segment the EC into the MEC and LEC homologues based on the connectivity with other regions, a voxel-by-voxel measure of connectivity probability was needed. We therefore also ran the tractography only seeding from the EC ROIs. Then, for each voxel in the ROI, we counted how many of the seeded paths reached the other ROIs. These connectivity counts were normalized to a probability, providing connectivity maps for the EC with the other four ROIs. The resulting smoothed and thresholded group averaged connectivity maps are shown in Figure 2. The sagittal slices show that the connectivity with presubiculum and RSC appears to be strongest in the posterior part of the EC, whereas the connectivity with dCA1pSub and OFC is strongest anteriorly in the EC. Further, the presubiculum connectivity does not show a clear medial-lateral gradient, but the connections with dCA1pSub, RSC and OFC are stronger laterally in the EC in the selected coronal slices.

**Figure 2:**
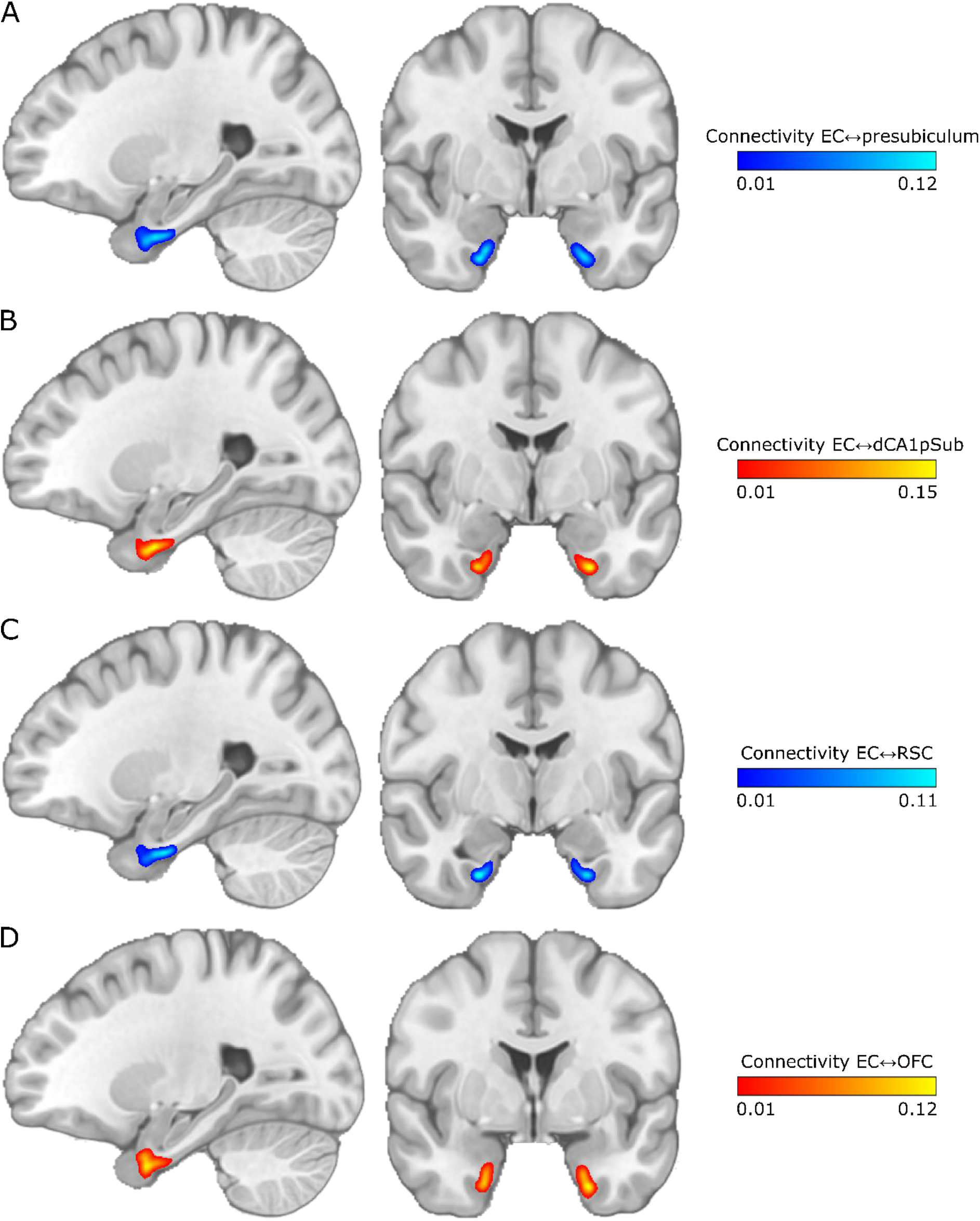
Group average maps of EC connectivity with EC and presubiculum, dCA1pSub, RSC and OFC. The maps are shown on sagittal (left) and coronal (right) slices in MNI space. The colormap intensity represents the fraction of paths seeded from that EC voxel that reached the other ROI. **A:** EC connectivity with presubiculum, **B:** EC connectivity with dCA1pSub, **C:** EC connectivity with RSC, **D:** EC connectivity with OFC.

Corresponding connectivity maps for one example participant are shown in Figure 2–figure supplement 1.

For segmentation into the MEC and LEC homologues, the main hypothesis was that these regions could be identified based on connectivity with presubiculum vs. dCA1pSub, respectively. The actual segmentation was performed on a voxel-by-voxel level in the EC determining with which of the other two regions the connection probability was highest, using the connectivity maps described in the previous paragraph. For comparison, the MEC-LEC segmentation was also performed based on connectivity with RSC vs. OFC, respectively. This was first performed individually for all participants, and inter-participant segmentation variability maps for the presubiculum vs. dCA1pSub and RSC vs. OFC segmentation approaches are shown in Figure 3. For most participants, MEC is clearly located more posteriorly and LEC is located more anteriorly for both segmentation approaches, and in addition they are located more medially and laterally with respect to each other for the presubiculum vs. dCA1pSub approach. The RSC vs. OFC approach also shows this medial-lateral trend of MEC and LEC across participants, although not as clear as for presubiculum vs. dCA1pSub. Corresponding MEC and LEC segmentations for one example participant are shown in Figure 3–figure supplement 1.

**Figure 3:**
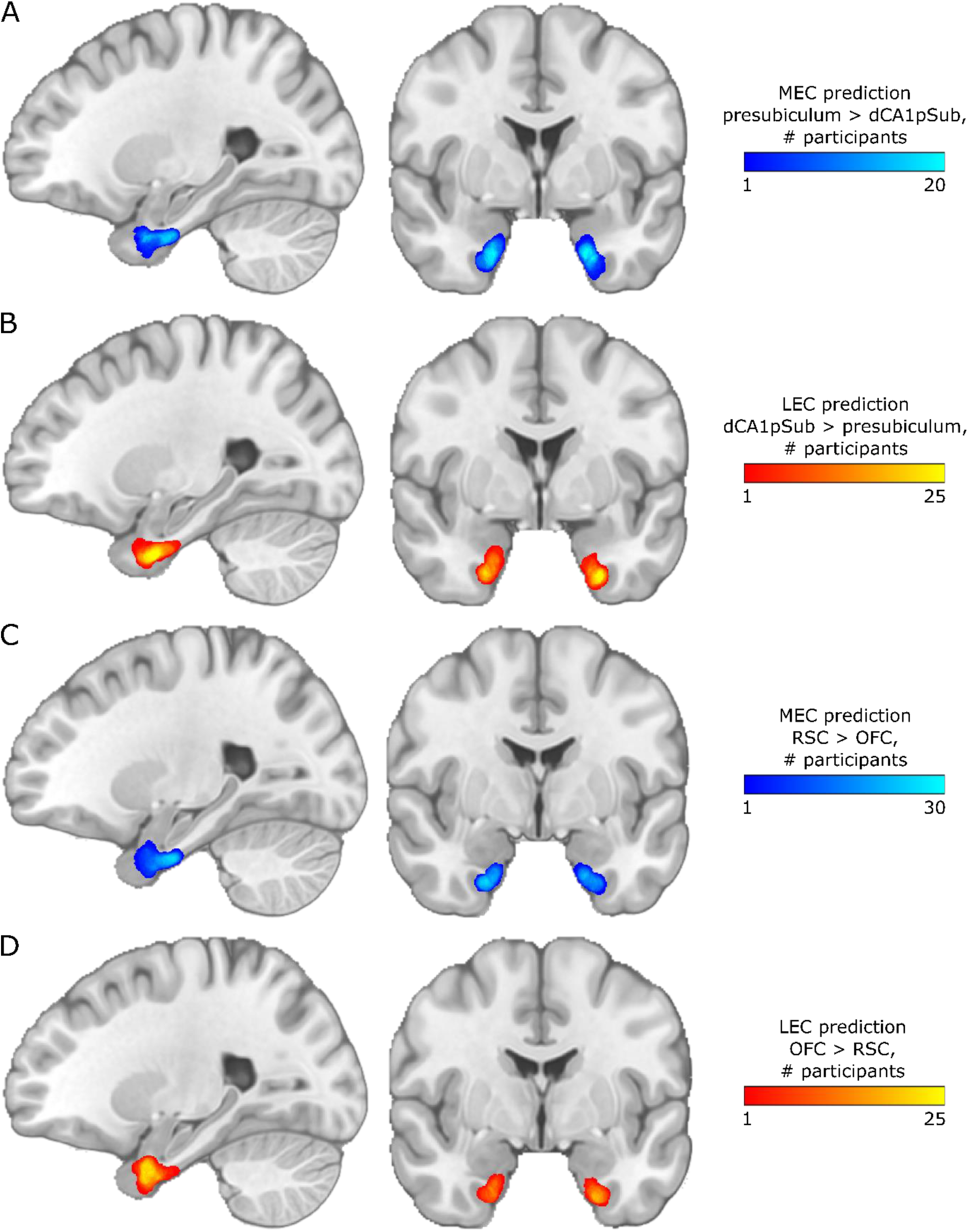
Inter-participant segmentation variability maps for different segmentation approaches. Results are shown on sagittal (left) and coronal (right) slices in MNI space. The colormap intensities represents the number of participants for which that voxel was classified as MEC or LEC, respectively. **A:** MEC prediction based on higher connectivity with presubiculum than with dCA1pSub, **B:** LEC prediction based higher connectivity with dCA1pSub than with presubiculum, **C:** MEC prediction based on higher connectivity with RSC than with OFC, **D:** LEC prediction based on higher connectivity with OFC than with RSC.

The same connectivity-based MEC-LEC segmentation was performed on a group level using the group averaged connectivity maps from Figure 2. As described above, the group segmentation was also performed using two different approaches – presubiculum vs. dCA1pSub, and RSC vs. OFC – and the resulting segmentations are shown in Figure 4. We see that for the MEC and LEC predictions from presubiculum vs. dCA1pSub, there is a clear medial-lateral (ML) and posterior-anterior (PA)-oriented border between the subregions. For RSC vs. OFC, however, the PA-oriented border is most prominent, but it is also slightly ML-oriented, most visible in the left EC. Because the results from the two approaches were slightly different, we also tried to interchange the order of the ROIs, and MEC and LEC segmentations from using presubiculum vs. OFC and RSC vs. dCA1pSub can be seen in Figure 4–figure supplement 1. Furthermore, to include all the information from the 2×2 combinations of seed regions into one final segmentation, we performed another approach where we averaged the connectivity maps for presubiculum and RSC, and the maps for dCA1pSub and OFC (Figure 5A and B). Figure 5C shows the resulting MEC and LEC homologues from this combined segmentation approach. With this approach, as with separate combinations of seed regions, we find both a PA- and ML-oriented (although most visible in the left hemisphere) border between MEC and LEC. These final MEC and LEC masks are also available in the Supplementary files.

**Figure 4:**
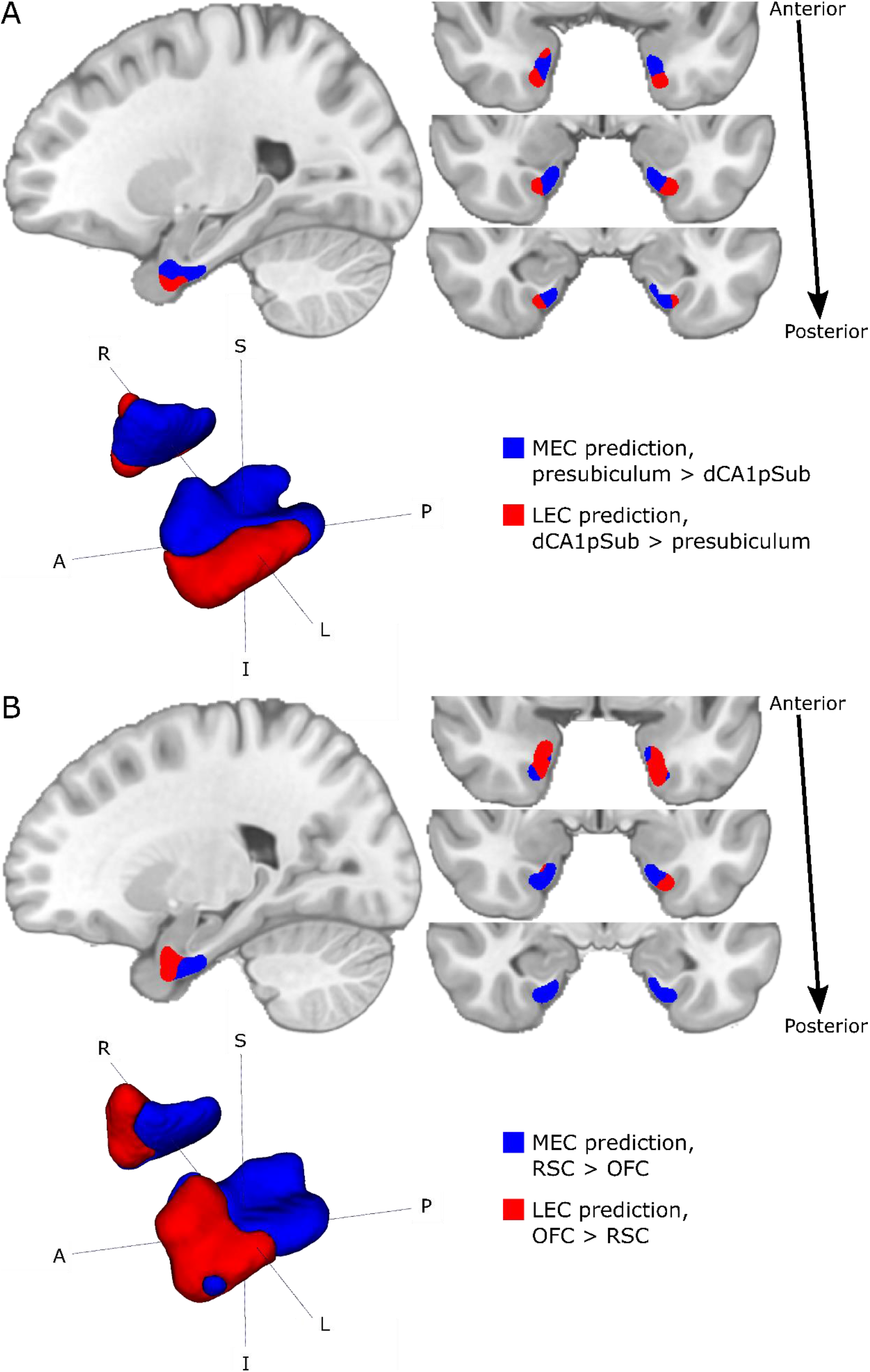
Group segmentations of MEC and LEC from different approaches. Results are shown on sagittal (top left) and coronal (top right) slices and 3D-rendered (bottom left) in MNI space. The MEC and LEC predictions are shown in blue and red, respectively. **A:** MEC and LEC prediction based on connectivity with presubiculum vs. dCA1pSub, **B:** MEC and LEC prediction based on connectivity with RSC vs. OFC. S = superior, I = inferior, A = anterior, P = posterior, R = right, L = left.

**Figure 5:**
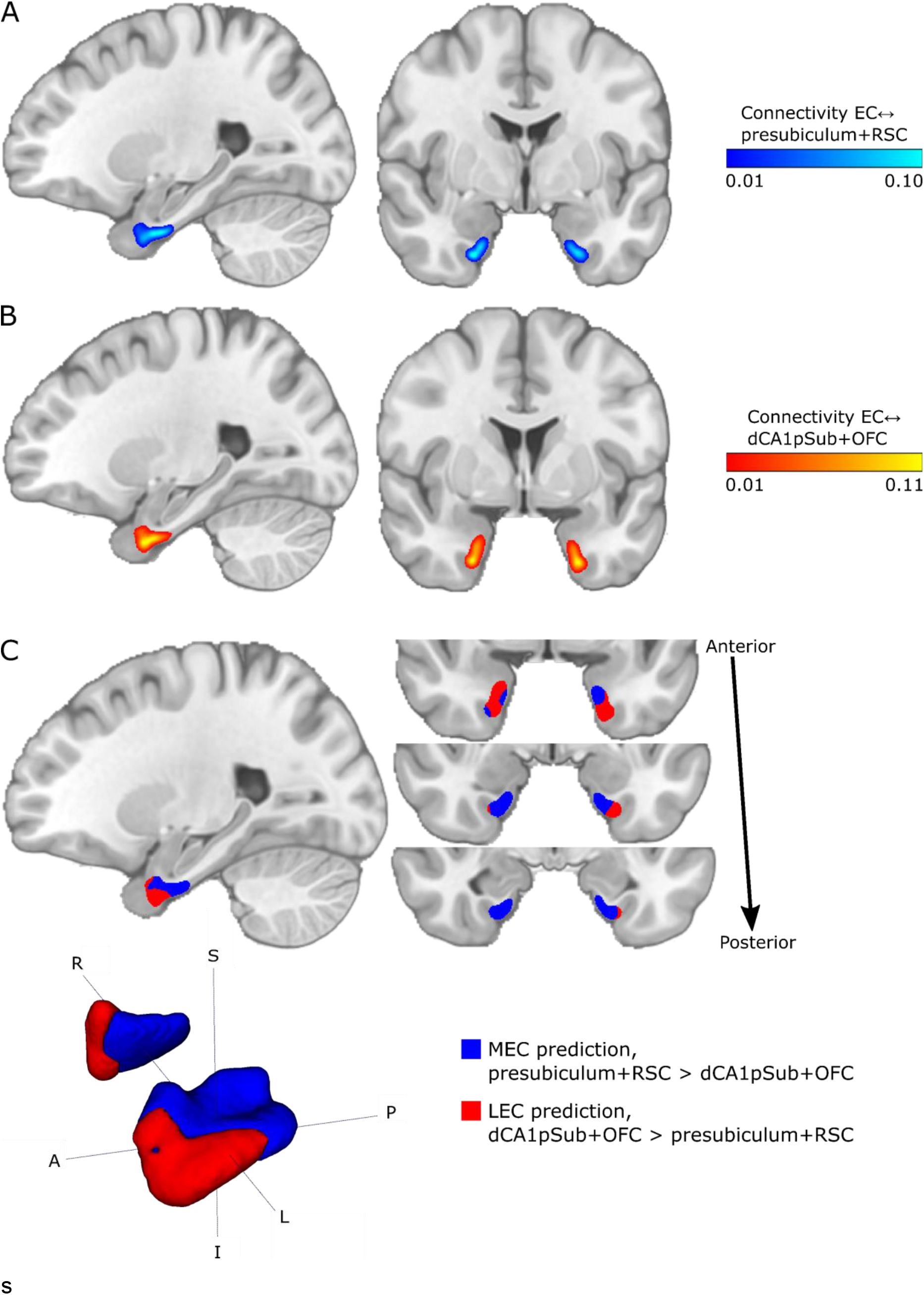
Group connectivity maps and segmentation using a combined approach with presubiculum + RSC vs. dCA1pSub + OFC. **A:** EC connectivity with presubiculum + RSC combined. **B:** EC connectivity with dCA1pSub + OFC combined. **C:** MEC and LEC prediction based on connectivity with presubiculum + RSC vs. dCA1pSub + OFC combined.

In a next step, since the borders of the segmentations from different approaches showed slightly different orientations along the posterior-anterior (PA) and medial-lateral (ML) axes, we wanted to quantify this directional difference by calculating the “degree” of PA- and ML-orientation of the borders. This was defined as a percentage from 0 to 100%, dependent on the angle between the MEC-LEC center of gravity vector and a pure PA or ML vector. Table 1 shows the resulting degrees of PA-vs. ML-oriented borders for the different segmentation approaches including the fMRI segmentations from previous studies (Maass et al., 2015; Navarro Schröder et al., 2015). The center of gravity vectors are also plotted in a common reference frame in Table 1–figure supplement 1. All DTI segmentation approaches have a border with a PA-orientation of around 50-60%, and a varying degree of ML-orientation from 6% for RSC vs. OFC up to 67% for presubiculum vs. dCA1pSub. The borders between the segmentations from fMRI have a high PA-orientation of around 92%, and a lower degree of ML-orientation than all of the DTI approaches. Interestingly, when comparing the different combinations of DTI approaches, using dCA1pSub as the defining region for LEC yields a higher degree of ML-orientation than using OFC. Similarly, using RSC as the defining region for MEC yields a slightly higher degree of PA-orientation of the border than using presubiculum, but this is less prominent.

**Table 1:** Degree of posterior-anterior (PA) or medial-lateral (ML)-orientation of the border between MEC and LEC for different segmentation approaches. The degree of PA- or ML-orientation is given as a percentage between 0 and 100%, dependent on the angle between the MEC-LEC center of gravity vector and the pure PA or ML vector, respectively. All numbers are given as the mean of both hemispheres ± mean absolute deviation.

| Segmentation approach |  | Posterior-anterior (PA) axis |  | Medial-lateral (ML) axis |  |
| --- | --- | --- | --- | --- | --- |
|  |  | Angle (°) | % PA | Angle (°) | % ML |
| DTI | Presubiculum/dCA1pSub | 45.1 ± 3.0 | 49.9 ± 3.3 | 29.8 ± 1.9 | 66.9 ± 2.1 |
| | RSC/OFC | 39.7 ± 15.3 | 55.9 ± 17.0 | 84.5 ± 11.1 | 6.1 ± 12.3 |
| | Presubiculum/OFC | 41.3 ± 17.3 | 54.1 ± 19.3 | 81.2 ± 10.3 | 9.8 ± 11.5 |
| | RSC/dCA1pSub | 33.9 ± 1.2 | 62.3 ± 1.3 | 64.9 ± 18.8 | 27.9 ± 20.9 |
| | Presubiculum+RSC/dCA1pSub+OFC | 37.5 ± 10.6 | 58.3 ± 11.8 | 73.1 ± 17.4 | 18.7 ± 19.3 |
| fMRI | Navarro Schröder et al. | 6.8 ± 1.8 | 92.4 ± 2.0 | 85.5 ± 0.7 | 5.0 ± 0.8 |
| | Maass et al. | 6.6 ± 0.5 | 92.7 ± 0.6 | 87.9 ± 0.9 | 2.4 ± 1.0 |

Finally, we wanted to compare the resulting sizes of the MEC and LEC homologues from all the different segmentation approaches, and these are shown in Table 2. For all DTI approaches, the MEC is larger than LEC, while fMRI on the other hand yields a larger LEC than MEC. The subregions are most equally sized when using the RSC vs. dCA1pSub approach.

**Table 2:** Resulting sizes of MEC and LEC for different segmentation approaches, and the size ratio between MEC and LEC. The numbers of voxels are given for the ROIs in MNI space with 0.5 mm isotropic resolution.

| Segmentation approach |  | Size (# voxels) |  | MEC/LEC<br>size ratio |
| --- | --- | --- | --- | --- |
|  |  | MEC | LEC |  |
| DTI | Presubiculum/dCA1pSub | 12759 | 7763 | 1.64 |
|  | RSC/OFC | 12971 | 8727 | 1.49 |
|  | Presubiculum/OFC | 13614 | 6979 | 1.95 |
|  | RSC/dCA1pSub | 11045 | 10282 | 1.07 |
|  | Presubiculum+RSC/dCA1pSub+OFC | 13571 | 7379 | 1.84 |
| fMRI | Navarro Schröder et al. | 12802 | 16028 | 0.80 |
|  | Maass et al. | 3776 | 11008 | 0.34 |

## Discussion

In this study, we used DTI and probabilistic tractography in 35 healthy adults to segment the human EC into homologues of what in other mammals have been functionally and cytoarchitectonically defined as MEC and LEC. We based the segmentation on EC connectivity with four brain regions known to selectively project to either of the EC subregions in multiple species. Different combinations of these four regions all showed both a posterior-anterior (PA) and a medial-lateral (ML)-oriented border between the human homologues of MEC and LEC. This orientation of the thus defined border is similar to that defined in previous fMRI studies resulting in the definition of the two subregions as pmEC and alEC (Maass et al., 2015; Navarro Schröder et al., 2015). Note however that our DTI results show a larger degree of ML-orientation, and a correspondingly lower degree of PA-orientation of the border between the subregions compared to the previous fMRI results.

The results from our study substantiate the pmEC and alEC subdivision of the human EC suggested in previous fMRI studies (Maass et al., 2015; Navarro Schröder et al., 2015). Although some earlier fMRI studies on mnemonic processing in the EC found a dissociation primarily along the medial-lateral axis (Reagh & Yassa, 2014; Schultz et al., 2012), it is important to realize that even the orientation of the cytoarchitectonically defined border between MEC and LEC in rodents does not align along a pure medial-to-lateral axis. Rather, the MEC in rodents is actually located in the posterior-medial EC, and the LEC is located in the anterior-lateral EC (van Strien et al., 2009). Also, in macaque monkeys, tracing studies show differential connectivity in caudal vs. rostral portions (Witter & Amaral, 2021). A pure medial-lateral subdivision of human EC is thus not to be expected. Nevertheless, the somewhat different orientations of the border between the human homologues of MEC vs. LEC subdivisions found using DTI vs. fMRI studies raises the question of which of the two imaging modalities should be preferred to define the position and orientation of this border.

There are several possible explanations as to why our DTI study showed slightly different segmentation results than the fMRI studies. First, DTI and fMRI are two different imaging modalities with inherently different mechanisms of connectivity. While DTI exploits the diffusion of water molecules in order to trace the structural paths of connectivity between brain regions (Mori et al., 1999; Mori & Zhang, 2006; Powell et al., 2004; Zeineh et al., 2012), fMRI identifies functional connectivity by correlating blood-oxygen-level-dependent (BOLD) signals across time (Van Dijk et al., 2010). Although structural and functional connectivity in theory should be closely linked, they are in reality quantitatively difficult to compare because of the complexity of the connectivity mechanisms of the brain (H. Huang & Ding, 2016; Messé, Benali, & Marrelec, 2015). Another reason for the different results between this and the previous studies could be the use of different seed regions to identify the MEC and LEC homologues. While we used presubiculum and RSC to define MEC, and dCA1pSub and OFC to define LEC (Caballero-Bleda & Witter, 1993; Honda & Ishizuka, 2004; Hoover & Vertes, 2007; Jones & Witter, 2007; Kondo & Witter, 2014; Saleem et al., 2008; Witter & Amaral, 1991; Witter & Amaral, 2021; Wyss & Van Groen, 1992), one of the fMRI studies investigated differential connectivity of PHC vs. PRC and distal vs. proximal subiculum (Maass et al., 2015), whereas the other used regions in a posterior-medial vs. an anterior-temporal cortical system (Navarro Schröder et al., 2015). The new insights from rodent anatomy indicate that while PHC area TH is connected with the MEC, PHC area TF is connected with the LEC (Witter & Amaral, 2021). As area TF is located more laterally than TH, this might in part explain why the previous fMRI study where they used connectivity with the whole PHC to define the pmEC (Maass et al., 2015) showed a lower medial-lateral component of their pmEC-alEC segmentation than our results. In order to determine to which extent each of these reasons contributed to the different subdivision results across studies, both imaging modalities and different seed regions should be investigated more rigorously in one single, larger cohort of participants.

Interestingly, using different seed regions to identify MEC and LEC resulted in varying degrees of PA- and ML-orientation of the border between them. It is unclear whether this is inherently linked to the DTI method, or due to an actual connectivity difference between the regions. Using presubiculum and dCA1pSub as the seed regions, which are situated medially and laterally with respect to each other, respectively, resulted in a border with higher degree of ML-than PA-orientation. On the other hand, using RSC and OFC, which are situated posteriorly and anteriorly in the brain, respectively, resulted in a border with higher degree of PA-than ML-orientation. Although it is not unnatural to assume that the brain is organized such that connected regions are situated more closely to each other, this could also be an effect of using probabilistic tractography, where the apparent connectivity probability depends on e.g. the length of the path and the size of the ROIs (Behrens, Johansen-Berg, Jbabdi, Rushworth, & Woolrich, 2007). In other species, including rodents and monkeys, the presubiculum and RSC show inputs to the EC with a similar spatial distribution (Witter & Amaral, 2021), aligning with our maps of connectivity paths with these two seed regions. However, comparing the different MEC and LEC segmentations from the different seed region combinations shows that while interchanging presubiculum and RSC yields only slightly different orientation of the border along the PA and ML axes, the difference when interchanging dCA1pSub and OFC is more substantial. In other species, dCA1pSub are known to project to both rostral and dorsolateral parts of EC, whereas posterolateral OFC mainly projects dorsolaterally in the EC (Kondo & Witter, 2014; Saleem et al., 2008; Witter & Amaral, 1991; Witter & Amaral, 2021). Whether these regions in humans project to different parts of the homologue of LEC, or whether our results are affected by using DTI and probabilistic tractography, should be further investigated by also comparing EC functional connectivity to these areas using fMRI. Note also that the location of projections from dCA1pSub along the medial-lateral axis of the EC depends on where the seed is placed along the posterior-anterior axis of the dCA1pSub (Witter & Amaral, 2021), which emphasizes the importance of anatomically accurate seed ROIs.

In order to determine and compare the connectivities between EC and the other ROIs, we normalized the connectivity maps by dividing them by the maximum probability of each map. This could introduce a bias in the results. By doing this, we intrinsically assume that the maximum connectivity strength to each of the other ROIs are equal, and the segmentation process does not take into account that the MEC connections might be stronger than the LEC connections, or vice versa. However, little is known about the strength of connectivities at this level of detail, particularly since it is not straightforward to examine or even define connectivity strength. Connectivity strength surely depends on axonal density, but other factors like synaptic density and efficacy are other important variables. Nevertheless, even if we were to know that some of the connections are stronger than the others, probabilistic tractography provides a relative instead of an absolute measure of connectivity and is also dependent on path lengths, ROI sizes and the number of possible path directions in a voxel. Normalizing the connectivity maps based on different connectivity strengths would therefore be a highly complex task. Therefore, we did not impose any further assumptions about connectivity strengths in our analyses.

Our study has some limitations. To define our ROIs, we chose to use regions from automatic cortical segmentation protocols. This could have influenced the anatomical precision of our analysis. Manual segmentation would be labor-intensive and requires high skills in neuroanatomy, possibly limiting the number of participants that could be included in the study. However, we manually adjusted some of the automatically segmented ROIs, and also intersected the registered ROIs from MNI space with the participants’ individual automatic segmentations in order to increase the anatomical accuracy. Another limitation is that there are inherent challenges to the EPI sequence used for diffusion imaging. This results in a generally low signal-to-noise ratio in the EC and the whole medial temporal lobe. In addition, these regions appear geometrically distorted in the EPI images, and although this has been corrected for, it is not possible to recover all of the lost signal. Imperfect correction can also affect the accuracy of the ROIs. Because of the probabilistic nature of the tractography technique it is unlikely that noise will introduce false connections, but it can leave some connections undetected. At last, a relatively low number of participants were included in our study, which might have influenced the statistical power of the results.

In conclusion, our DTI results support the definition of pmEC and alEC as the human homologues of MEC and LEC. Also inspired by novel insights coming from rodent anatomy, we present a segmentation based on a combination of differential presubiculum/RSC and dCA1pSub/OFC structural connectivity which indicates a border between the two subdivisions of EC with an orientation that is angled both towards the posterior-anterior axis, as well as to the medial-lateral axis. The fact that there are some differences in the orientation of the border based on DTI and fMRI data in addition to the seed regions used, indicates the need for investigation in a larger number of participants across both modalities.

## Materials and methods

### MRI data

Structural and diffusion MRI data from 35 healthy adults were obtained from the MGH-USC Human Connectome Project (https://ida.loni.usc.edu, http://db.humanconnectome.org) (Fan et al., 2016). The data were acquired on a Siemens 3T Connectom scanner with maximum gradient strength of 300 mT/m and slew rate 200 T/m/s (McNab et al., 2013; Setsompop et al., 2013). Structural T1-weighted images were acquired using a 3D magnetization-prepared rapid gradient-echo (MPRAGE) sequence at 1 mm isotropic resolution. Diffusion data were acquired using a spin-echo echo-planar imaging (EPI) sequence at 1.5 mm isotropic resolution, with b-values of 1000 s/mm^2^ (64 directions), 3000 s/mm^2^ (64 directions), 5000 s/mm^2^ (128 directions) and 10,000 s/mm^2^ (256 directions). One non-diffusion-weighted (b = 0) image was collected every 14 image volumes.

### Preprocessing

The MRI data were minimally preprocessed by the Human Connectome Project as described in (Fan et al., 2014). In brief, this preprocessing pipeline included gradient nonlinearity correction, motion correction, Eddy current correction and b-vector correction.

#### Registration

Both structural and diffusion images were brain extracted using the brain mask from running the FreeSurfer functions *recon-all* and *dt-recon* on the participant’s structural and diffusion images, respectively (Fischl et al., 2002; Fischl et al., 2004), before refining the result using FSL’s BET (http://fsl.fmrib.ox.ac.uk/fsl/) (Jenkinson, Beckmann, Behrens, Woolrich, & Smith, 2012; Smith, 2002). For the diffusion images, brain extraction and registration were performed on the participant’s average b = 1000 image. The individual brain-extracted structural and diffusion images were registered to each other, as well as to the MNI152-09b standard brain template (Fonov, Evans, McKinstry, Almli, & Collins, 2009), using symmetric non-linear registration in the Advanced Neuroimaging Toolbox (ANTs) based on mutual information (Avants et al., 2011).

#### Regions of interest

Regions of interest (ROIs) including the EC, presubiculum, CA1 and subiculum were extracted from the automated cortical and subcortical parcellation obtained from running FreeSurfer’s *recon-all* and *segmentHA_T1* functions on the MNI152-09b template (Fischl et al., 2002; Fischl et al., 2004; Iglesias et al., 2015). The EC ROI was further refined by masking it by a probabilistic EC ROI, thresholded at 0.25 from the Jülich-Brain Cytoarchitectonic Atlas (Amunts, Mohlberg, Bludau, & Zilles, 2020). Since the resulting EC ROI extended too far posteriorly towards the parahippocampal cortex and laterally beyond the collateral sulcus, we also performed a manual adjustment. We created ROIs of distal CA1/proximal subiculum by splitting each of the two hippocampal structures in half along its proximodistal axis. Of all voxels encompassing CA1, the half located distally was included, and of all the voxels encompassing subiculum, the half located proximally was included: these two halves thus make up what we here define and refer to as ‘distal CA1/proximal subiculum’ (dCA1pSub). To create RSC and OFC ROIs, respectively, the FreeSurfer parcellations named “isthmus cingulate” and “lateral orbitofrontal” were used as a starting point. The final RSC ROI was obtained by tailoring the isthmus cingulate and removing the excess superior areas, while the final OFC ROI was obtained by extracting the posterolateral quadrant of the lateral orbitofrontal area. All resulting ROIs are shown in Figure 1–figure supplements 2–6. The ROIs were registered to the participants’ individual spaces by applying the calculated transformations from ANTs. To increase the anatomical precision of the ROIs, the registered ROIs were then masked by respective participant-specific FreeSurfer parcellations.

### DTI analysis

All DTI analyses were performed in the participant’s native diffusion space. Voxel-wise fiber orientation distribution functions (fODFs) were computed by running the FSL function *bedpostx* on the diffusion data, using the zeppelin deconvolution model, a Rician noise model, and burn-in period 3000 (Sotiropoulos et al., 2016). Probabilistic tractography between the EC and presubiculum, dCA1pSub, RSC and OFC ROIs was then performed by running FSL’s *probtrackx2* on the fODFs (Behrens et al., 2007; Behrens, Woolrich, et al., 2003). Tractography was performed both in ROI-by-ROI and voxel-by-ROI connectivity mode, with number of samples 250,000, minimal path length 5 mm, and a midline termination mask (Behrens, Johansen-Berg, et al., 2003; Ezra et al., 2015; Johansen-Berg et al., 2004; Máté et al., 2018; Saygin et al., 2011). For tractography between EC and presubiculum, paths were excluded if they reached the dCA1pSub ROI, while for tractography between EC and dCA1pSub, paths were excluded if they reached the presubiculum ROI – and equivalently for tractography between EC and RSC/OFC. For both *bedpostx* and *probtrackx2,* parameters were run with default values unless otherwise specified. ROI-by-ROI connectivity mode provides probability maps of the connectivity paths between the ROIs, while voxel-by-ROI connectivity mode provides probability maps of the voxel-wise connectivity of the EC ROI with the other ROIs, respectively. All tractography results were registered to MNI space and further analyses were performed there to facilitate inter-participant comparisons.

### MEC and LEC segmentation

The voxel-wise connectivity maps were normalized to [0,1] by dividing by the maximum probability for each hemisphere separately, and then thresholded by 0.01 to reduce false positive connections (Behrens, Johansen-Berg, et al., 2003; Saygin et al., 2011). This threshold was determined empirically by testing a range of thresholds and choosing the one that in most cases removed connections outside the grey matter, because due to remaining distortions in the DTI images some of the EC ROIs slightly extended into white matter and air voxels. Crucially, we then define the MEC as the region that is most strongly connected with the presubiculum and/or RSC, while the LEC is the region that is most strongly connected with dCA1pSub and/or OFC (Caballero-Bleda & Witter, 1993; Honda & Ishizuka, 2004; Hoover & Vertes, 2007; Jones & Witter, 2007; Kondo & Witter, 2014; Saleem et al., 2008; Witter & Amaral, 1991; Witter & Amaral, 2021; Wyss & Van Groen, 1992). For each participant, a hard segmentation was performed on the normalized and thresholded voxel-wise connectivity maps using FSL’s *find_the_biggest* (Behrens, Johansen-Berg, et al., 2003; Johansen-Berg et al., 2004), where the voxels that had a stronger connection probability with the presubiculum/RSC than with dCA1pSub/OFC were classified as MEC, and vice versa for LEC.

### Group analysis

Group probability maps of the connectivity paths between the ROIs, as well as group probability maps of voxel-wise connectivity, were created by summing and averaging all the individual maps. Inter-participant segmentation variability maps were created by adding together all the individual participants’ MEC and LEC segmentations, respectively. Group MEC and LEC segmentation were performed similarly to the individual segmentation: The group voxel-wise connectivity maps were first smoothed with a Gaussian kernel of 1 mm and thresholded by 0.01, and then a hard segmentation was performed equivalently to the single-participant segmentation by comparing the connection probabilities of EC with presubiculum/RSC vs. dCA1pSub/OFC. Four different segmentations were performed with all the 2×2 combinations of seed regions, in addition to a combined segmentation approach where the connectivity maps for presubiculum + RSC and for dCA1pSub + OFC, respectively, were combined and averaged before segmentation.

### Segmentation comparisons

To assess the different segmentation approaches and compare the resulting locations of MEC and LEC, we calculated the orientation of the MEC-LEC border along the posterior-anterior (PA) and medial-lateral (ML) axes, respectively. This was performed by first calculating the centers of gravity of the differently defined MECs and LECs, and the vector between these centers of gravity. Next, the angle between this vector and a pure PA or ML vector was determined. The degree of PA- or ML-oriented border was then defined between 0 and 100% such that an angle of 0° to the PA or ML vector means that the border is 100% oriented along the PA or ML vector, respectively. Correspondingly, an angle of 90° would mean that the border is 0% oriented along the respective axis, i.e. it is orthogonal to that axis. In addition, the different segmentations were compared with respect to the sizes of the resulting MECs and LECs, and the size ratios between these were calculated. All these segmentation comparisons were also carried out on the two fMRI-based segmentations of pmEC and alEC available for download from earlier studies (Maass et al., 2015; Navarro Schröder et al., 2015).

## Acknowledgements

Data were provided by the Human Connectome Project, MGH-USC Consortium (Principal Investigators: Bruce R. Rosen, Arthur W. Toga and Van Wedeen; U01MH093765) funded by the NIH Blueprint Initiative for Neuroscience Research grant; the National Institutes of Health grant P41EB015896; and the Instrumentation Grants S10RR023043, 1S10RR023401, 1S10RR019307.

## Supplementary files

- MEC and LEC homologue masks in MNI152-09b space, defined from the combined presubiculum+RSC vs. dCA1pSub+OFC approach (MEC_LEC_segmentations.zip).

---

**Figure 1–figure supplement 1:**
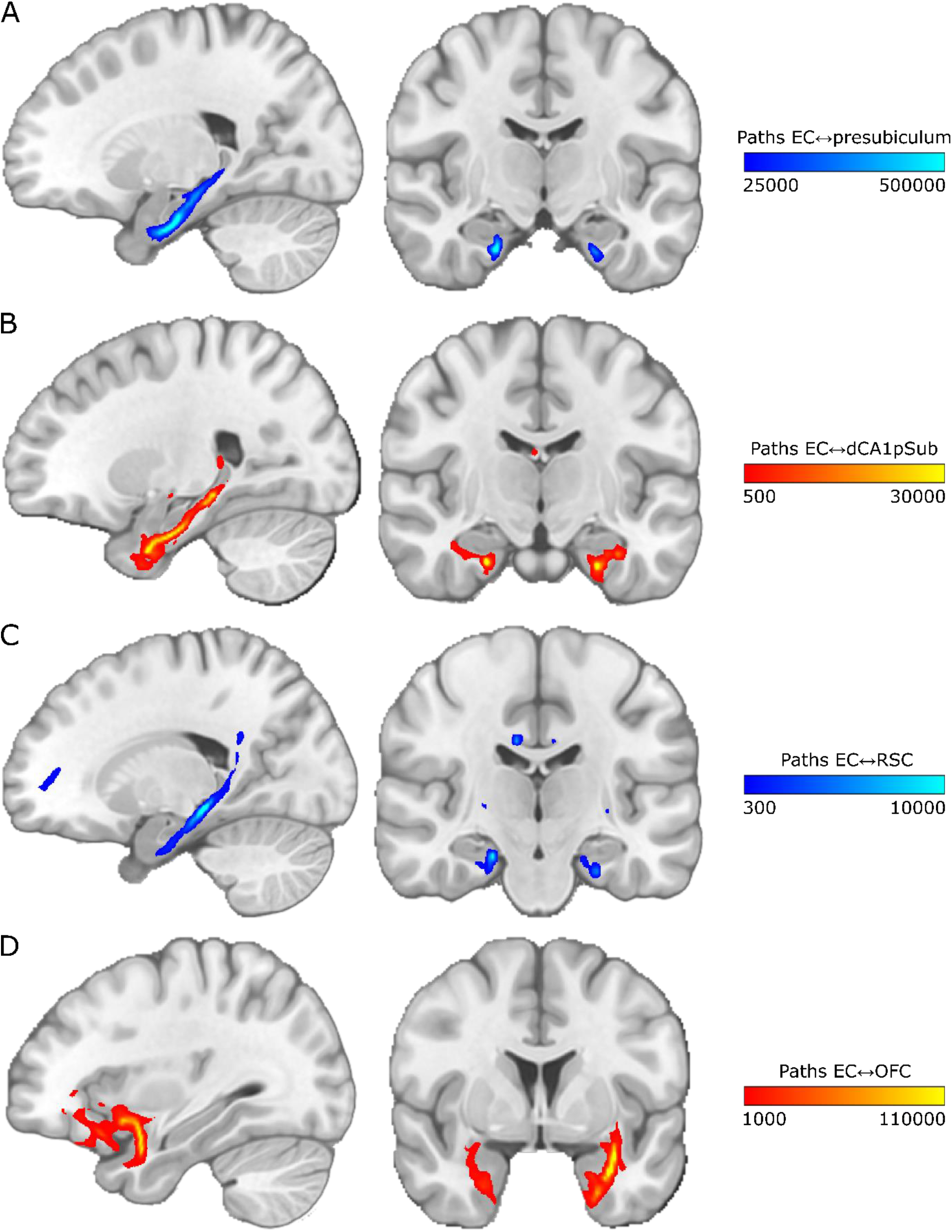
Connectivity paths between EC and presubiculum, dCA1pSub, RSC and OFC for one example participant. The paths are shown on sagittal (left) and coronal (right) slices in MNI space. The colormap intensity represents the number of probabilistic paths running through that voxel. **A:** Paths between EC and presubiculum, **B:** Paths between EC and dCA1pSub, **C:** Paths between EC and RSC, **D:** Paths between EC and OFC.

**Figure 1–figure supplement 2:**
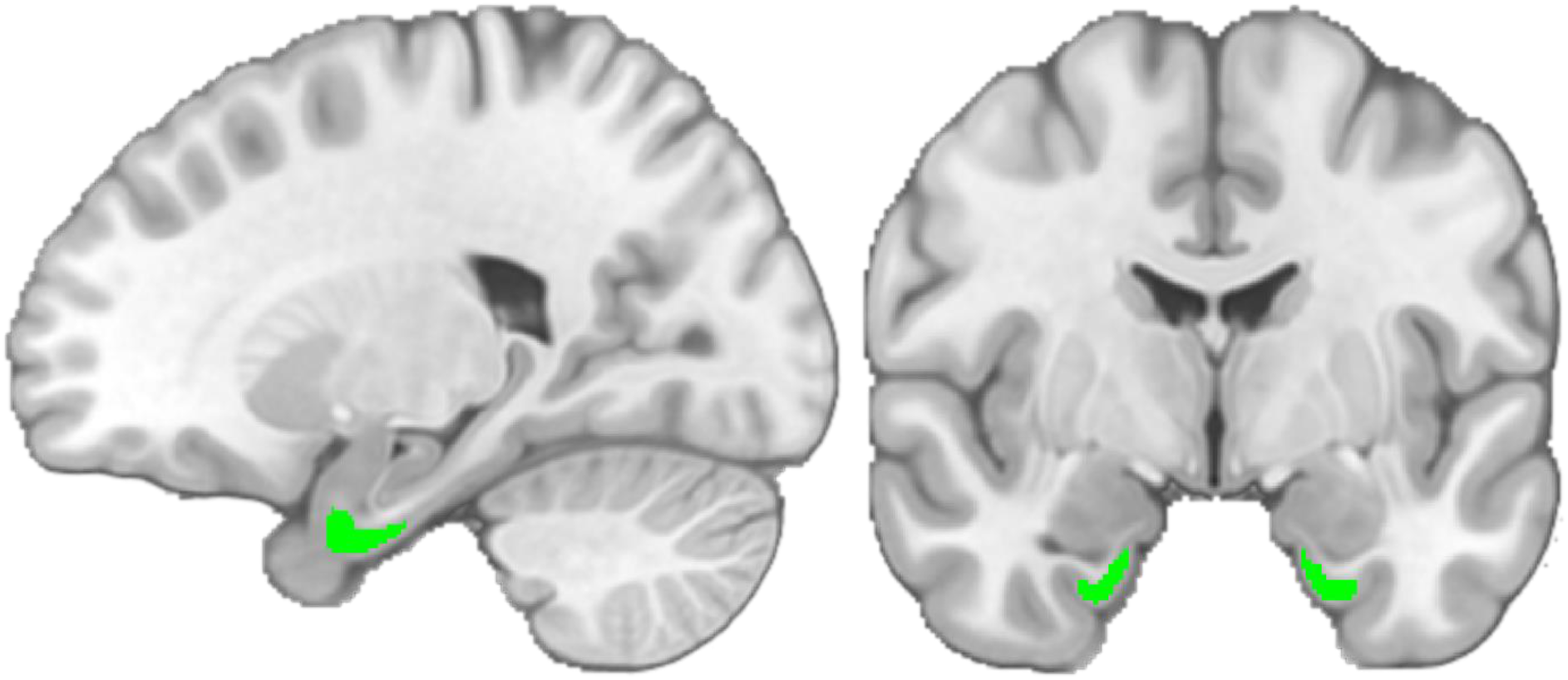
EC ROI. The ROI is shown in green on a sagittal (left) and a coronal (right) slice in MNI space.

**Figure 1–figure supplement 3:**
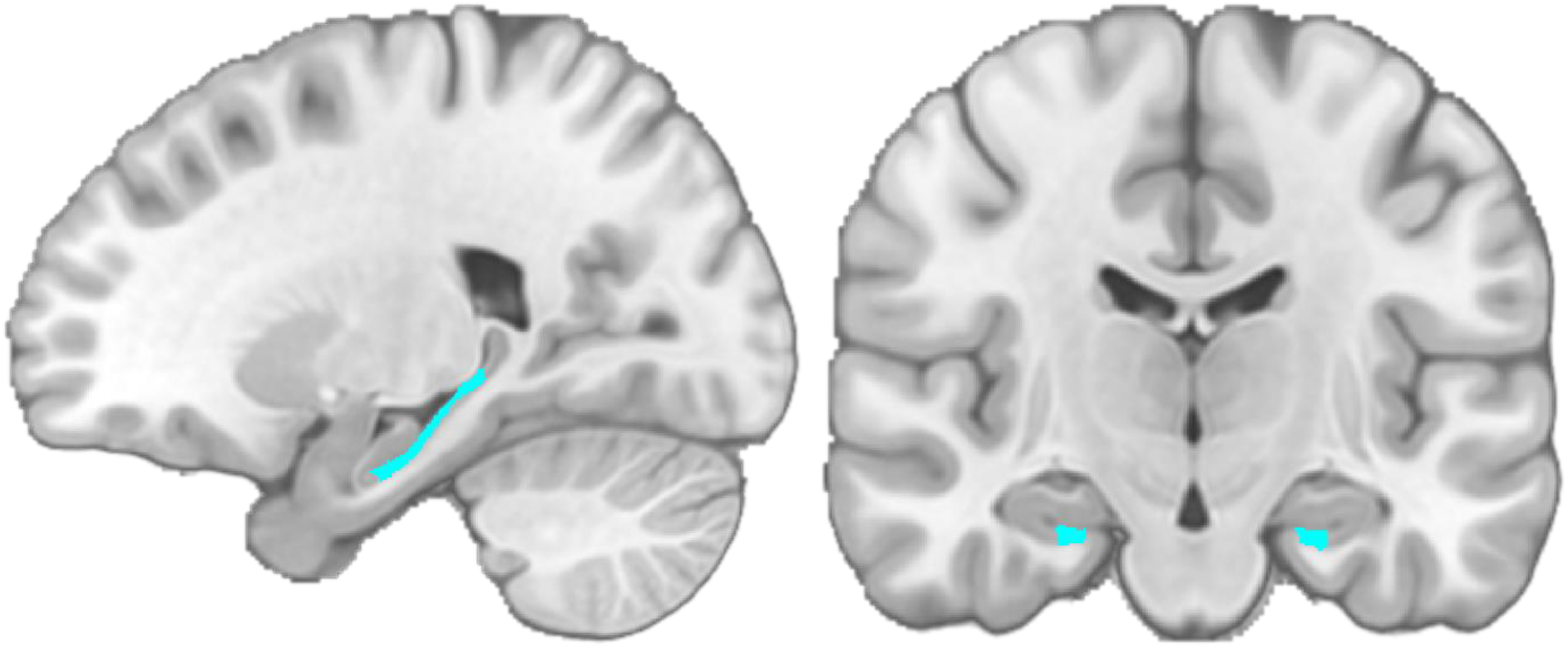
Presubiculum ROI. The ROI is shown in light blue on a sagittal (left) and a coronal (right) slice in MNI space.

**Figure 1–figure supplement 4:**
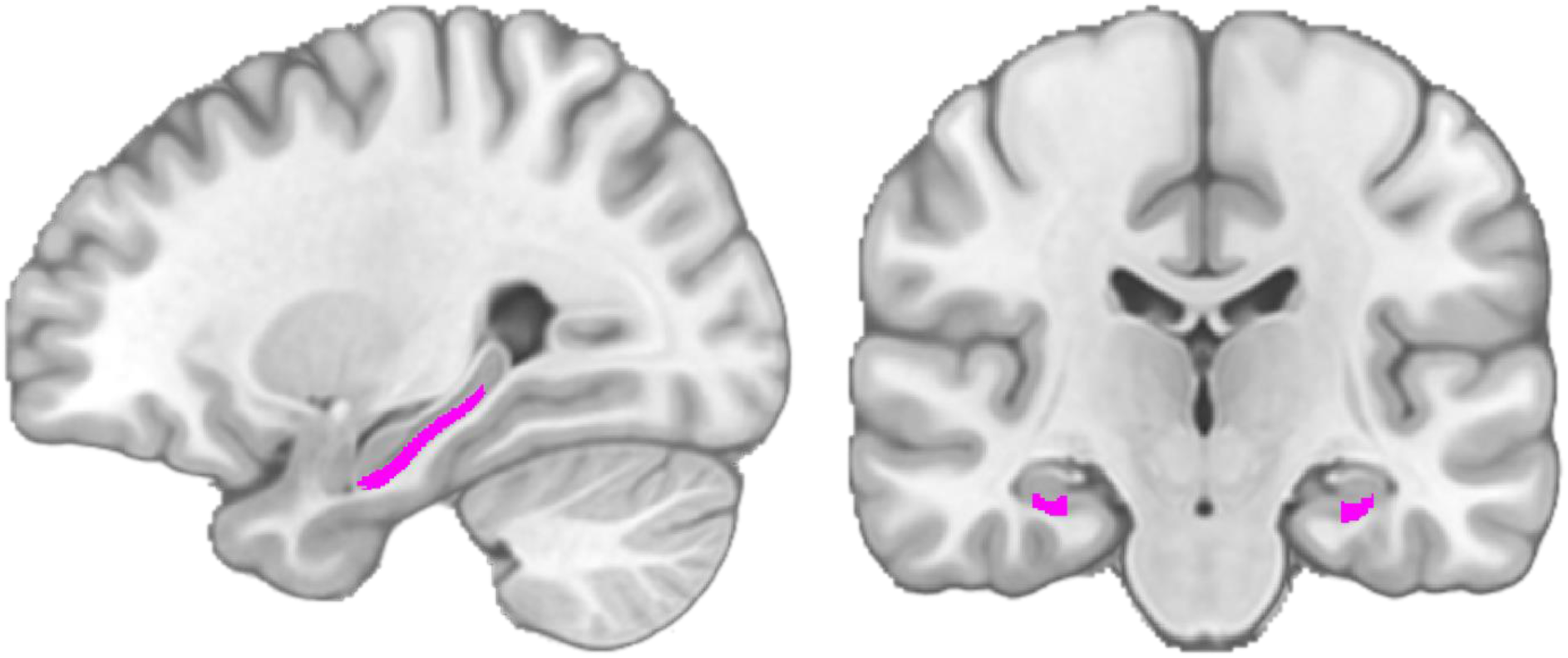
dCA1pSub ROI. The ROI is shown in pink on a sagittal (left) and a coronal (right) slice in MNI space.

**Figure 1–figure supplement 5:**
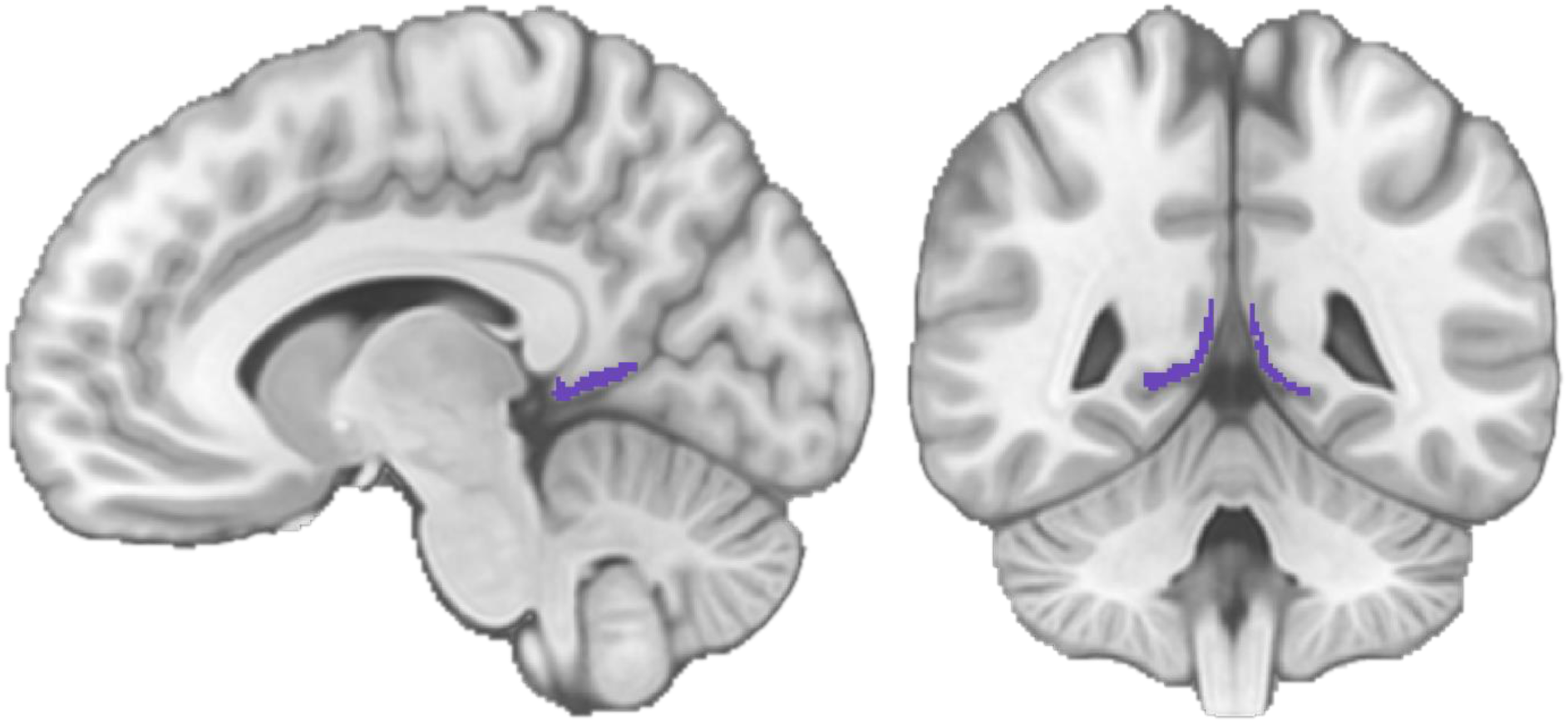
RSC ROI. The ROI is shown in purple on a sagittal (left) and a coronal (right) slice in MNI space.

**Figure 1–figure supplement 6:**
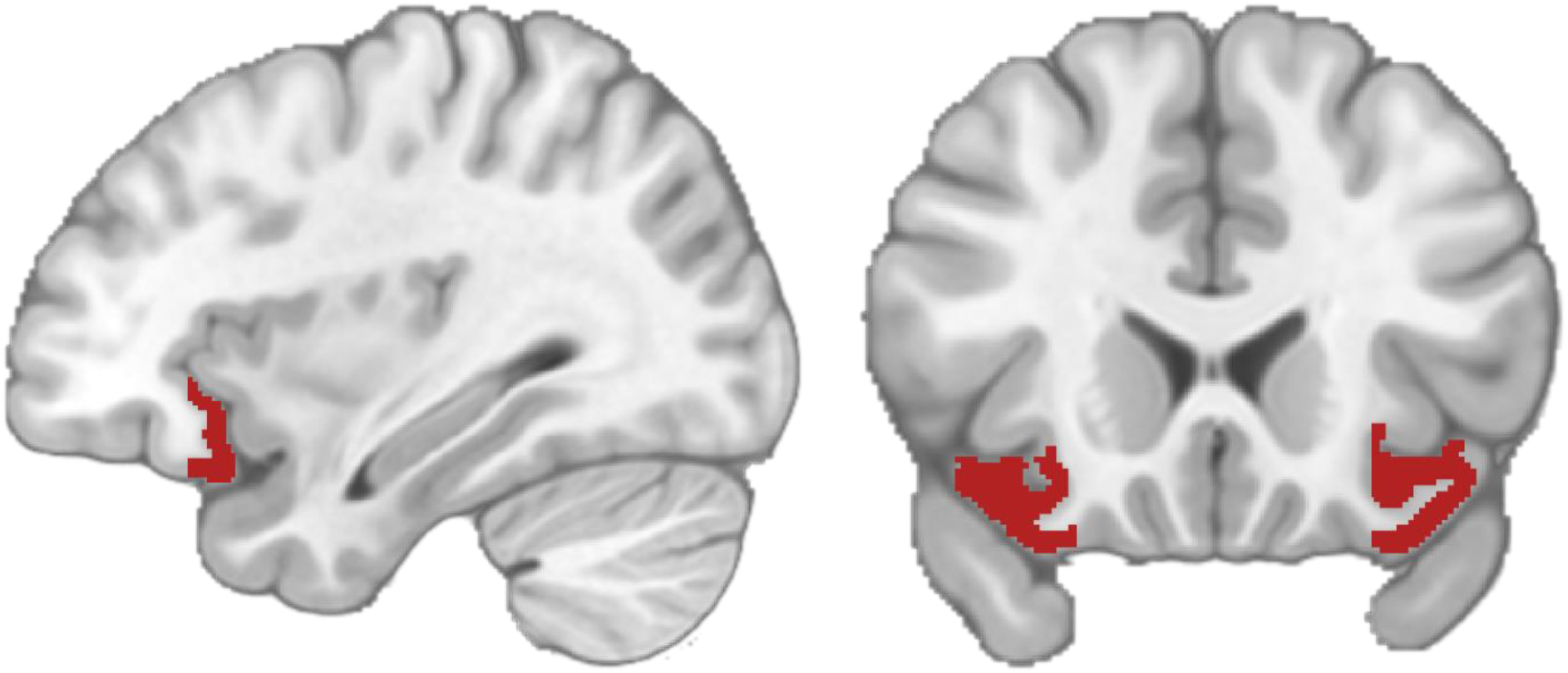
OFC ROI. The ROI is shown in dark red on a sagittal (left) and a coronal (right) slice in MNI space.

**Figure 2–figure supplement 1:**
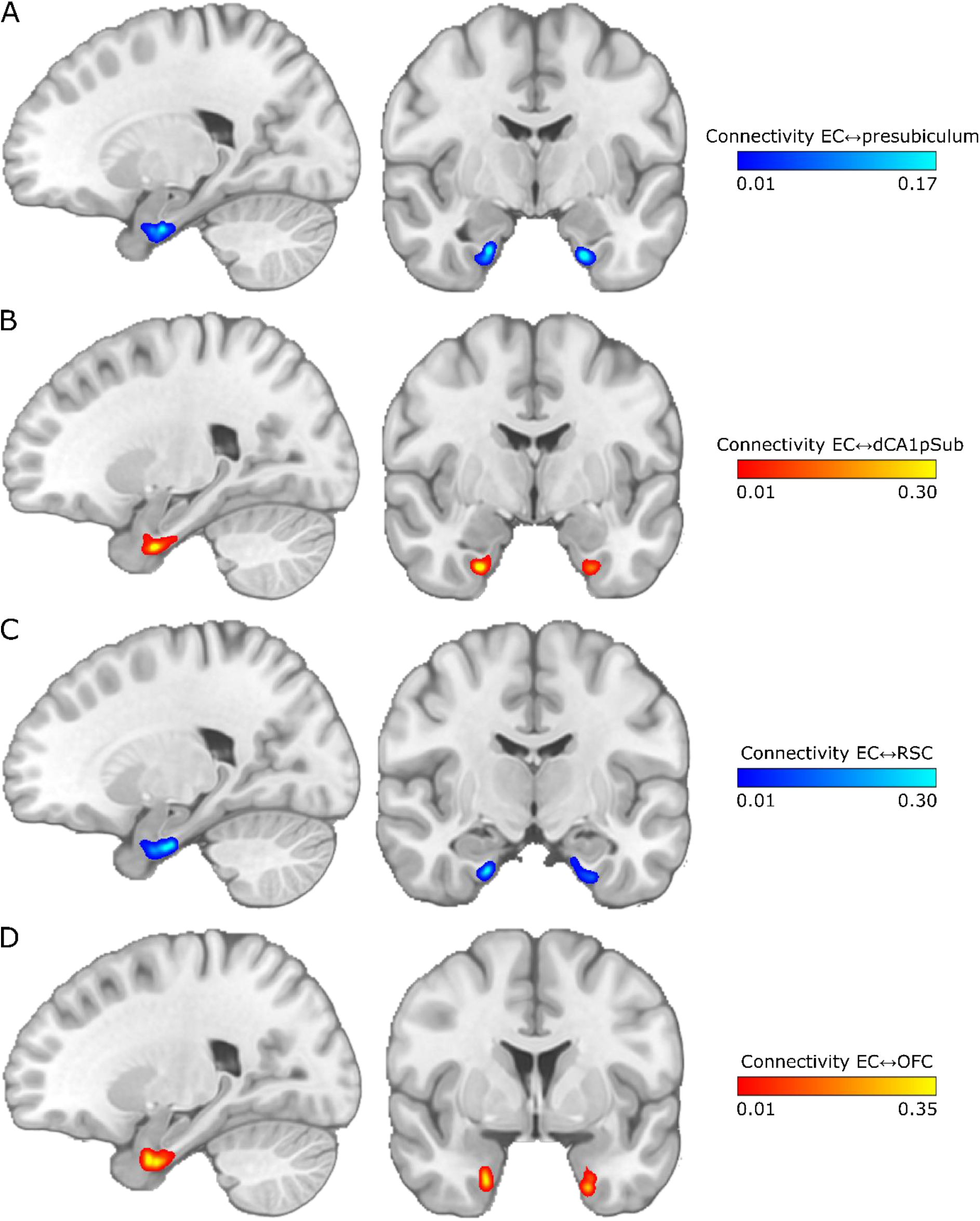
Maps of EC connectivity with presubiculum, dCA1pSub, RSC and OFC for one example participant. The maps are shown on sagittal (left) and coronal (right) slices in MNI space. The colormap intensity represents the fraction of paths seeded from that EC voxel that reached the other ROI. **A:** EC connectivity with presubiculum, **B:** EC connectivity with dCA1pSub, **C:** EC connectivity with RSC, **D:** EC connectivity with OFC.

**Figure 3–figure supplement 1:**
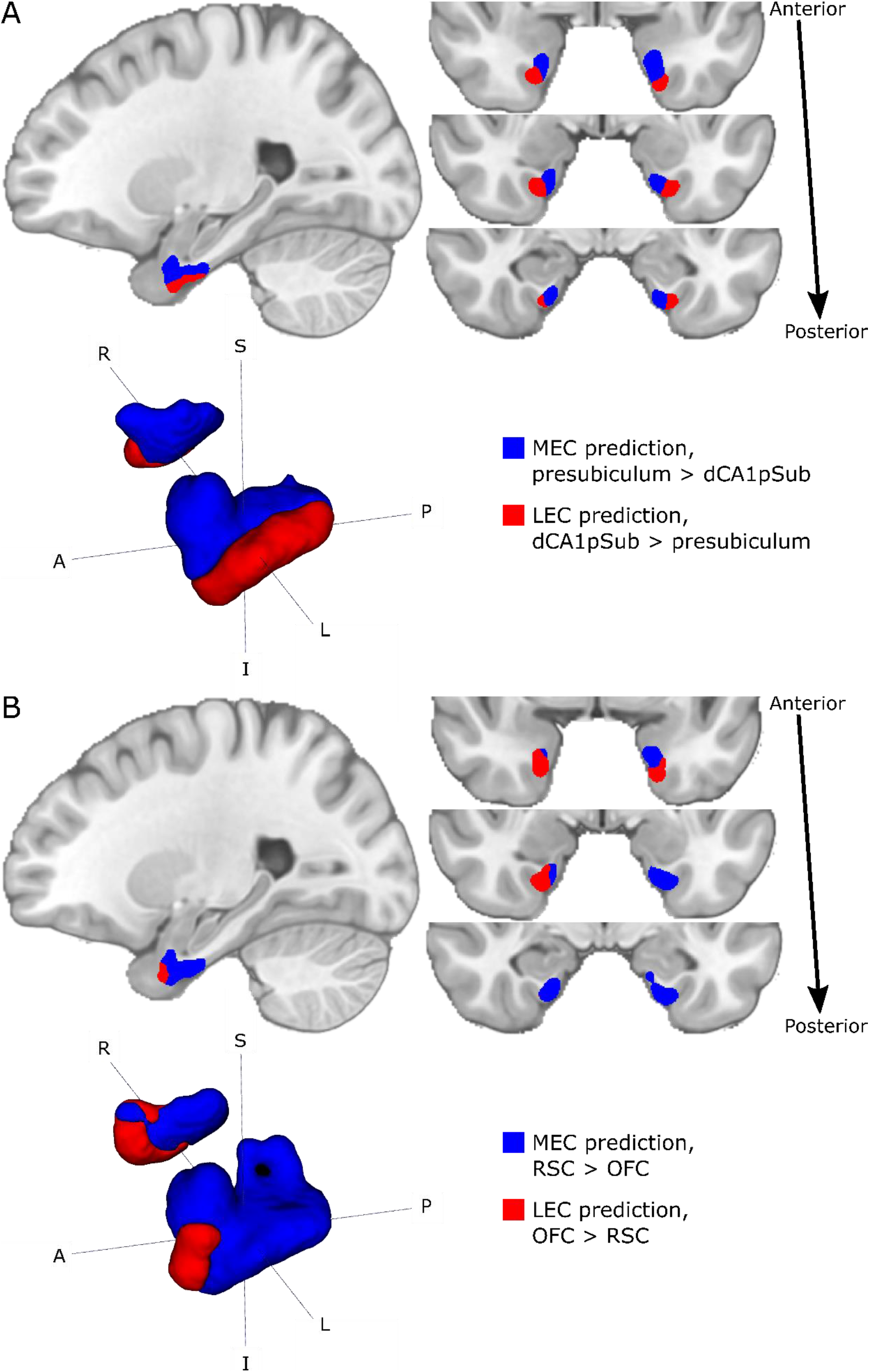
Segmentations of MEC and LEC from different approaches for one example participant. The results are shown on sagittal (top left) and coronal (top right) slices and 3D-rendered (bottom left) in MNI space. The MEC and LEC predictions are shown in blue and red, respectively. **A:** MEC and LEC prediction based on connectivity with presubiculum vs. dCA1pSub, **B:** MEC and LEC prediction based on connectivity with RSC vs. OFC. S = superior, I = inferior, A = anterior, P = posterior, R = right, L = left.

**Figure 4–figure supplement 1:**
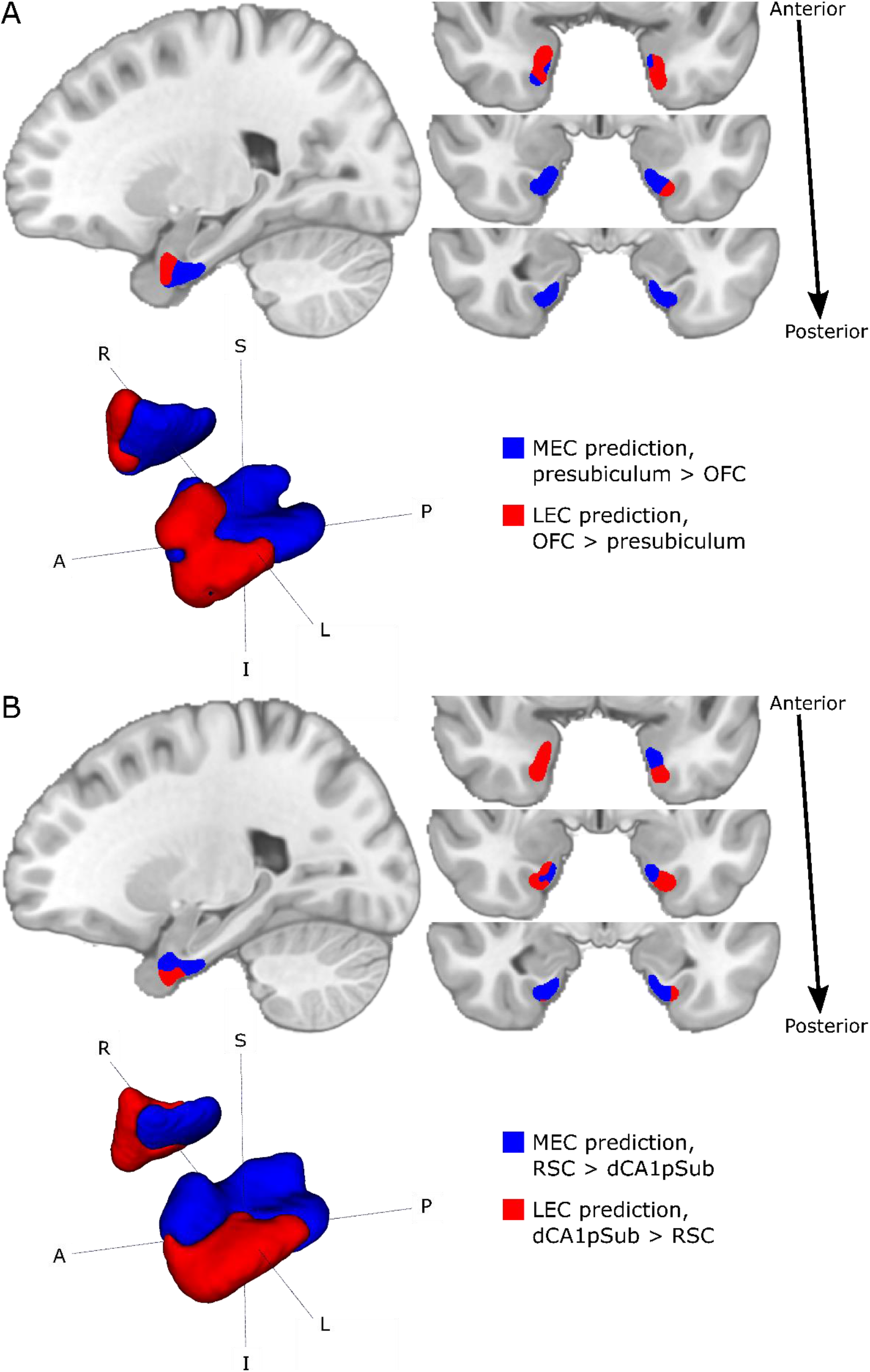
Group segmentations of MEC and LEC from different approaches. The results are shown on sagittal (top left) and coronal (top right) slices and 3D-rendered (bottom left) in MNI space. The MEC and LEC predictions are shown in blue and red, respectively. **A:** MEC and LEC prediction based on connectivity with presubiculum vs. OFC, **B:** MEC and LEC prediction based on connectivity with RSC vs. dCA1pSub. S = superior, I = inferior, A = anterior, P = posterior, R = right, L = left.

**Table 1–figure supplement 1:**
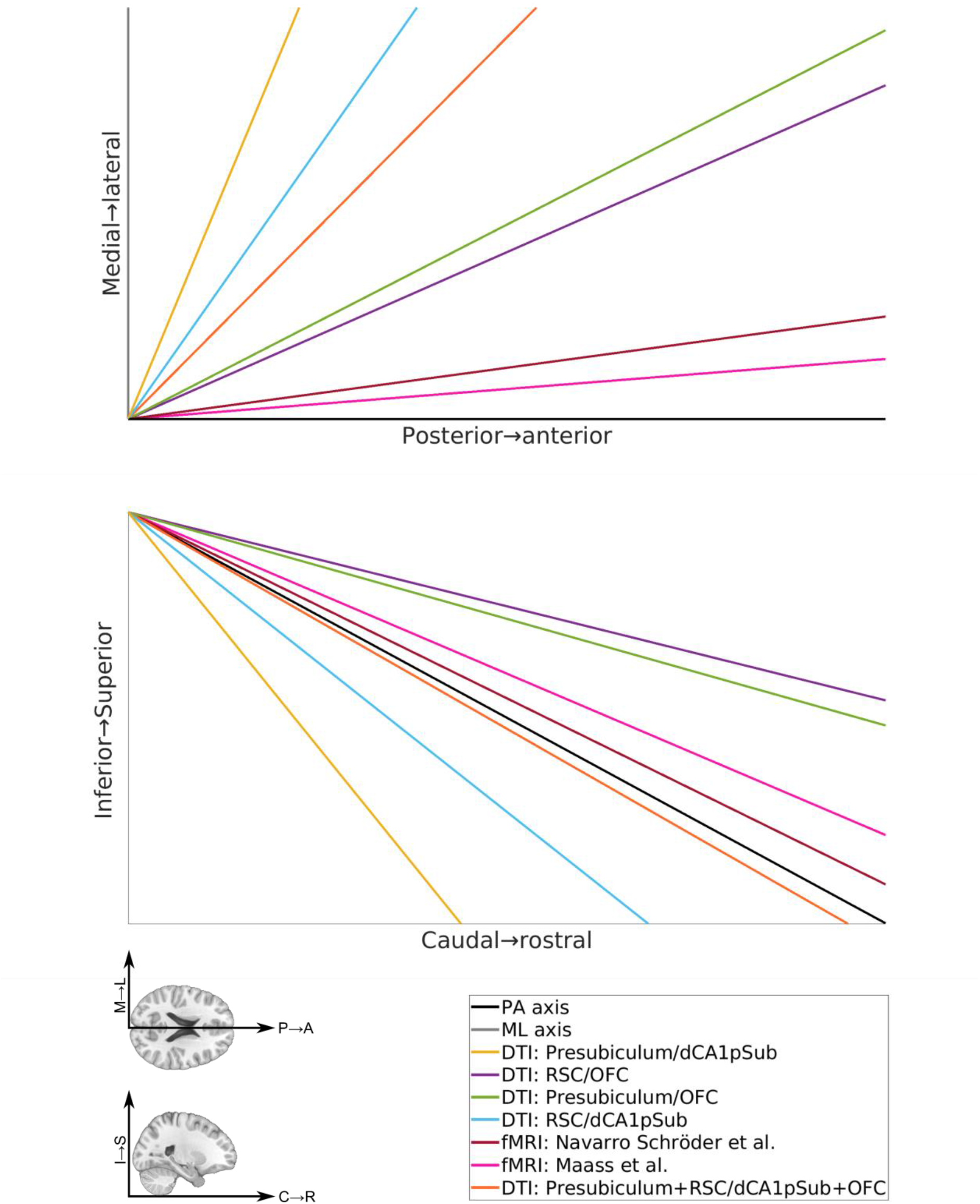
Center of gravity vectors between MEC-LEC segmentations in the axial (top) and sagittal (bottom) planes, showing the angles between the orientation of the MEC-LEC border and the pure PA and ML axes in the left hemisphere. The PA axis vector is shown in black, the ML axis vector is shown in grey (visible in the axial view only), and the colors of the MEC-LEC vectors for all the different segmentation approaches are explained in the legend box on the bottom right. On the bottom left, illustrations of the anatomical directions of the vector plots are shown (M = medial, L = lateral, P = posterior, A = anterior, I = interior, S = superior, C = caudal, R = rostral). The posterior-anterior axis is defined along the long axis of the hippocampus in the sagittal plane. Note that the length of the vectors shown here does not correspond to the real distance between the MEC and LEC centers of gravity. The origin of the vectors corresponds to the MEC center of gravity, which are not the true MEC locations for the different segmentation approaches, but shifted in space so that all vectors originate in the same point.

## References

Amunts, K., Mohlberg, H., Bludau, S., & Zilles, K. (2020). Julich-Brain: A 3D probabilistic atlas of the human brain’s cytoarchitecture. Science, 369(6506), 988–992. doi:10.1126/science.abb4588

Avants, B. B., Tustison, N. J., Song, G., Cook, P. A., Klein, A., & Gee, J. C. (2011). A reproducible evaluation of ANTs similarity metric performance in brain image registration. Neuroimage, 54(3), 2033–2044. doi:10.1016/j.neuroimage.2010.09.025

Behrens, T. E., Johansen-Berg, H., Jbabdi, S., Rushworth, M. F., & Woolrich, M. W. (2007). Probabilistic diffusion tractography with multiple fibre orientations: What can we gain? Neuroimage, 34(1), 144–155. doi:10.1016/j.neuroimage.2006.09.018

Behrens, T. E., Johansen-Berg, H., Woolrich, M. W., Smith, S. M., Wheeler-Kingshott, C. A., Boulby, P. A., Barker, G. J., Sillery, E. L., Sheehan, K., Ciccarelli, O., Thompson, A. J., Brady, J. M., & Matthews, P. M. (2003). Non-invasive mapping of connections between human thalamus and cortex using diffusion imaging. Nature Neuroscience, 6(7), 750–757. doi:10.1038/nn1075

Behrens, T. E., Woolrich, M. W., Jenkinson, M., Johansen-Berg, H., Nunes, R. G., Clare, S., Matthews, P. M., Brady, J. M., & Smith, S. M. (2003). Characterization and propagation of uncertainty in diffusion-weighted MR imaging. Magnetic Resonance in Medicine, 50(5), 1077–1088. doi:10.1002/mrm.10609

Bellmund, J. L., Deuker, L., & Doeller, C. F. (2019). Mapping sequence structure in the human lateral entorhinal cortex. eLife, 8. doi:10.7554/eLife.45333

Braak, H., & Braak, E. (1992). The human entorhinal cortex: normal morphology and lamina-specific pathology in various diseases. Neuroscience Research, 15(1), 6–31. doi:10.1016/0168-0102(92)90014-4

Buzsáki, G. (1996). The Hippocampo-Neocortical Dialogue. Cerebral Cortex, 6(2), 81–92. doi:10.1093/cercor/6.2.81

Caballero-Bleda, M., & Witter, M. P. (1993). Regional and laminar organization of projections from the presubiculum and parasubiculum to the entorhinal cortex: an anterograde tracing study in the rat. Journal of Comparative Neurology, 328(1), 115–129. doi:10.1002/cne.903280109

Canto, C. B., Wouterlood, F. G., & Witter, M. P. (2008). What Does the Anatomical Organization of the Entorhinal Cortex Tell Us? Neural Plasticity, 2008, 381243. doi:10.1155/2008/381243

Chen, X., Vieweg, P., & Wolbers, T. (2019). Computing distance information from landmarks and self-motion cues - Differential contributions of anterior-lateral vs. posterior-medial entorhinal cortex in humans. Neuroimage, 202, 116074. doi:10.1016/j.neuroimage.2019.116074

Deshmukh, S. S., & Knierim, J. J. (2011). Representation of non-spatial and spatial information in the lateral entorhinal cortex. Frontiers in Behavioral Neuroscience, 5, 69. doi:10.3389/fnbeh.2011.00069

Doan, T. P., Lagartos-Donate, M. J., Nilssen, E. S., Ohara, S., & Witter, M. P. (2019). Convergent Projections from Perirhinal and Postrhinal Cortices Suggest a Multisensory Nature of Lateral, but Not Medial, Entorhinal Cortex. Cell Reports, 29(3), 617–627.e617. doi:10.1016/j.celrep.2019.09.005

Doeller, C. F., Barry, C., & Burgess, N. (2010). Evidence for grid cells in a human memory network. Nature, 463(7281), 657–661. doi:10.1038/nature08704

Eichenbaum, H., Yonelinas, A. P., & Ranganath, C. (2007). The Medial Temporal Lobe and Recognition Memory. Annual Review of Neuroscience, 30(1), 123–152. doi:10.1146/annurev.neuro.30.051606.094328

Ezra, M., Faull, O. K., Jbabdi, S., & Pattinson, K. T. (2015). Connectivity-based segmentation of the periaqueductal gray matter in human with brainstem optimized diffusion MRI. Human Brain Mapping, 36(9), 3459–3471. doi:10.1002/hbm.22855

Fan, Q., Nummenmaa, A., Witzel, T., Zanzonico, R., Keil, B., Cauley, S., Polimeni, J. R., Tisdall, D., Van Dijk, K. R., Buckner, R. L., Wedeen, V. J., Rosen, B. R., & Wald, L. L. (2014). Investigating the capability to resolve complex white matter structures with high b-value diffusion magnetic resonance imaging on the MGH-USC Connectom scanner. Brain Connectivity, 4(9), 718–726. doi:10.1089/brain.2014.0305

Fan, Q., Witzel, T., Nummenmaa, A., Van Dijk, K. R. A., Van Horn, J. D., Drews, M. K., Somerville, L. H., Sheridan, M. A., Santillana, R. M., Snyder, J., Hedden, T., Shaw, E. E., Hollinshead, M. O., Renvall, V., Zanzonico, R., Keil, B., Cauley, S., Polimeni, J. R., Tisdall, D., Buckner, R. L., Wedeen, V. J., Wald, L. L., Toga, A. W., & Rosen, B. R. (2016). MGH-USC Human Connectome Project datasets with ultra-high b-value diffusion MRI. Neuroimage, 124(Pt B), 1108–1114. doi:10.1016/j.neuroimage.2015.08.075

Fischl, B., Salat, D. H., Busa, E., Albert, M., Dieterich, M., Haselgrove, C., van der Kouwe, A., Killiany, R., Kennedy, D., Klaveness, S., Montillo, A., Makris, N., Rosen, B., & Dale, A. M. (2002). Whole brain segmentation: automated labeling of neuroanatomical structures in the human brain. Neuron, 33(3), 341–355. doi:10.1016/s0896-6273(02)00569-x

Fischl, B., van der Kouwe, A., Destrieux, C., Halgren, E., Ségonne, F., Salat, D. H., Busa, E., Seidman, L. J., Goldstein, J., Kennedy, D., Caviness, V., Makris, N., Rosen, B., & Dale, A. M. (2004). Automatically parcellating the human cerebral cortex. Cerebral Cortex, 14(1), 11-22. doi:10.1093/cercor/bhg087

Fonov, V. S., Evans, A. C., McKinstry, R. C., Almli, C. R., & Collins, D. L. (2009). Unbiased nonlinear average age-appropriate brain templates from birth to adulthood. Neuroimage, 47, S102. doi:10.1016/S1053-8119(09)70884-5

Fyhn, M., Molden, S., Witter, M. P., Moser, E. I., & Moser, M.-B. (2004). Spatial Representation in the Entorhinal Cortex. Science, 305(5688), 1258–1264. doi:10.1126/science.1099901

Hafting, T., Fyhn, M., Molden, S., Moser, M. B., & Moser, E. I. (2005). Microstructure of a spatial map in the entorhinal cortex. Nature, 436(7052), 801–806. doi:10.1038/nature03721

Honda, Y., & Ishizuka, N. (2004). Organization of connectivity of the rat presubiculum: I. Efferent projections to the medial entorhinal cortex. Journal of Comparative Neurology, 473(4), 463–484. doi:10.1002/cne.20093

Hoover, W. B., & Vertes, R. P. (2007). Anatomical analysis of afferent projections to the medial prefrontal cortex in the rat. Brain Structure & Function, 212(2), 149–179. doi:10.1007/s00429-007-0150-4

Huang, C.-C., Rolls, E. T., Hsu, C.-C. H., Feng, J., & Lin, C.-P. (2021). Extensive Cortical Connectivity of the Human Hippocampal Memory System: Beyond the “What” and “Where” Dual Stream Model. Cerebral Cortex. doi:10.1093/cercor/bhab113

Huang, H., & Ding, M. (2016). Linking Functional Connectivity and Structural Connectivity Quantitatively: A Comparison of Methods. Brain Connectivity, 6(2), 99–108. doi:10.1089/brain.2015.0382

Høydal, Ø. A., Skytøen, E. R., Andersson, S. O., Moser, M. B., & Moser, E. I. (2019). Object-vector coding in the medial entorhinal cortex. Nature, 568(7752), 400–404. doi:10.1038/s41586-019-1077-7

Iglesias, J. E., Augustinack, J. C., Nguyen, K., Player, C. M., Player, A., Wright, M., Roy, N., Frosch, M. P., McKee, A. C., Wald, L. L., Fischl, B., & Van Leemput, K. (2015). A computational atlas of the hippocampal formation using ex vivo, ultra-high resolution MRI: Application to adaptive segmentation of in vivo MRI. Neuroimage, 115, 117–137. doi:10.1016/j.neuroimage.2015.04.042

Jenkinson, M., Beckmann, C. F., Behrens, T. E., Woolrich, M. W., & Smith, S. M. (2012). FSL. Neuroimage, 62(2), 782–790. doi:10.1016/j.neuroimage.2011.09.015

Johansen-Berg, H., Behrens, T. E., Robson, M. D., Drobnjak, I., Rushworth, M. F., Brady, J. M., Smith, S. M., Higham, D. J., & Matthews, P. M. (2004). Changes in connectivity profiles define functionally distinct regions in human medial frontal cortex. Proceedings of the National Academy of Sciences of the United States of America, 101(36), 13335–13340. doi:10.1073/pnas.0403743101

Jones, B. F., & Witter, M. P. (2007). Cingulate cortex projections to the parahippocampal region and hippocampal formation in the rat. Hippocampus, 17(10), 957–976. doi:10.1002/hipo.20330

Kerr, K. M., Agster, K. L., Furtak, S. C., & Burwell, R. D. (2007). Functional neuroanatomy of the parahippocampal region: The lateral and medial entorhinal areas. Hippocampus, 17(9), 697–708. doi:10.1002/hipo.20315

Knierim, J. J., Neunuebel, J. P., & Deshmukh, S. S. (2014). Functional correlates of the lateral and medial entorhinal cortex: objects, path integration and local-global reference frames. Philosophical Transactions of the Royal Society of London. Series B: Biological Sciences, 369(1635), 20130369. doi:10.1098/rstb.2013.0369

Kondo, H., & Witter, M. P. (2014). Topographic organization of orbitofrontal projections to the parahippocampal region in rats. Journal of Comparative Neurology, 522(4), 772–793. doi:10.1002/cne.23442

Lavenex, P., & Amaral, D. G. (2000). Hippocampal-neocortical interaction: A hierarchy of associativity. Hippocampus, 10(4), 420–430. doi:10.1002/1098-1063(2000)10:4<420::Aid-hipo8>3.0.Co;2-5

Máté, A., Kis, D., Czigner, A., Fischer, T., Halász, L., & Barzó, P. (2018). Connectivity-based segmentation of the brainstem by probabilistic tractography. Brain Research, 1690, 74–88. doi:10.1016/j.brainres.2018.03.010

McNab, J. A., Edlow, B. L., Witzel, T., Huang, S. Y., Bhat, H., Heberlein, K., Feiweier, T., Liu, K., Keil, B., Cohen-Adad, J., Tisdall, M. D., Folkerth, R. D., Kinney, H. C., & Wald, L. L. (2013). The Human Connectome Project and beyond: initial applications of 300 mT/m gradients. Neuroimage, 80, 234–245. doi:10.1016/j.neuroimage.2013.05.074

Messé, A., Benali, H., & Marrelec, G. (2015). Relating structural and functional connectivity in MRI: a simple model for a complex brain. IEEE Transactions on Medical Imaging, 34(1), 27–37. doi:10.1109/tmi.2014.2341732

Montchal, M. E., Reagh, Z. M., & Yassa, M. A. (2019). Precise temporal memories are supported by the lateral entorhinal cortex in humans. Nature Neuroscience, 22(2), 284–288. doi:10.1038/s41593-018-0303-1

Mori, S., Crain, B. J., Chacko, V. P., & van Zijl, P. C. (1999). Three-dimensional tracking of axonal projections in the brain by magnetic resonance imaging. Annals of Neurology, 45(2), 265–269. doi:10.1002/1531-8249(199902)45:2<265::aid-ana21>3.0.co;2-3

Mori, S., & Zhang, J. (2006). Principles of diffusion tensor imaging and its applications to basic neuroscience research. Neuron, 51(5), 527–539. doi:10.1016/j.neuron.2006.08.012

Moser, Edvard I., & Moser, M.-B. (2013). Grid Cells and Neural Coding in High-End Cortices. Neuron, 80(3), 765–774. doi:10.1016/j.neuron.2013.09.043

Maass, A., Berron, D., Libby, L. A., Ranganath, C., & Düzel, E. (2015). Functional subregions of the human entorhinal cortex. eLife, 4, e06426. doi:10.7554/eLife.06426

Navarro Schröder, T., Haak, K. V., Zaragoza Jimenez, N. I., Beckmann, C. F., & Doeller, C. F. (2015). Functional topography of the human entorhinal cortex. eLife, 4, e06738. doi:10.7554/eLife.06738

Nilssen, E. S., Doan, T. P., Nigro, M. J., Ohara, S., & Witter, M. P. (2019). Neurons and networks in the entorhinal cortex: A reappraisal of the lateral and medial entorhinal subdivisions mediating parallel cortical pathways. Hippocampus, 29(12), 1238–1254. doi:10.1002/hipo.23145

Powell, H. W., Guye, M., Parker, G. J., Symms, M. R., Boulby, P., Koepp, M. J., Barker, G. J., & Duncan, J. S. (2004). Noninvasive in vivo demonstration of the connections of the human parahippocampal gyrus. Neuroimage, 22(2), 740–747. doi:10.1016/j.neuroimage.2004.01.011

Ranganath, C., & Ritchey, M. (2012). Two cortical systems for memory-guided behaviour. Nature Reviews: Neuroscience, 13(10), 713–726. doi:10.1038/nrn3338

Reagh, Z. M., & Yassa, M. A. (2014). Object and spatial mnemonic interference differentially engage lateral and medial entorhinal cortex in humans. Proceedings of the National Academy of Sciences, 111(40), E4264–E4273. doi:10.1073/pnas.1411250111

Saleem, K. S., Kondo, H., & Price, J. L. (2008). Complementary circuits connecting the orbital and medial prefrontal networks with the temporal, insular, and opercular cortex in the macaque monkey. Journal of Comparative Neurology, 506(4), 659–693. doi:10.1002/cne.21577

Saygin, Z. M., Osher, D. E., Augustinack, J., Fischl, B., & Gabrieli, J. D. (2011). Connectivity-based segmentation of human amygdala nuclei using probabilistic tractography. Neuroimage, 56(3), 1353–1361. doi:10.1016/j.neuroimage.2011.03.006

Schultz, H., Sommer, T., & Peters, J. (2012). Direct Evidence for Domain-Sensitive Functional Subregions in Human Entorhinal Cortex. The Journal of Neuroscience, 32(14), 4716–4723. doi:10.1523/jneurosci.5126-11.2012

Setsompop, K., Kimmlingen, R., Eberlein, E., Witzel, T., Cohen-Adad, J., McNab, J. A., Keil, B., Tisdall, M. D., Hoecht, P., Dietz, P., Cauley, S. F., Tountcheva, V., Matschl, V., Lenz, V. H., Heberlein, K., Potthast, A., Thein, H., Van Horn, J., Toga, A., Schmitt, F., Lehne, D., Rosen, B. R., Wedeen, V., & Wald, L. L. (2013). Pushing the limits of in vivo diffusion MRI for the Human Connectome Project. Neuroimage, 80, 220–233. doi:10.1016/j.neuroimage.2013.05.078

Smith, S. M. (2002). Fast robust automated brain extraction. Human Brain Mapping, 17(3), 143–155. doi:10.1002/hbm.10062

Sotiropoulos, S. N., Hernández-Fernández, M., Vu, A. T., Andersson, J. L., Moeller, S., Yacoub, E., Lenglet, C., Ugurbil, K., Behrens, T. E. J., & Jbabdi, S. (2016). Fusion in diffusion MRI for improved fibre orientation estimation: An application to the 3T and 7T data of the Human Connectome Project. Neuroimage, 134, 396–409. doi:10.1016/j.neuroimage.2016.04.014

Suzuki, W. A., & Eichenbaum, H. (2000). The Neurophysiology of Memory. Annals of the New York Academy of Sciences, 911(1), 175–191. doi:10.1111/j.1749-6632.2000.tb06726.x

Tsao, A., Moser, M. B., & Moser, E. I. (2013). Traces of experience in the lateral entorhinal cortex. Current Biology, 23(5), 399–405. doi:10.1016/j.cub.2013.01.036

Tsao, A., Sugar, J., Lu, L., Wang, C., Knierim, J. J., Moser, M.-B., & Moser, E. I. (2018). Integrating time from experience in the lateral entorhinal cortex. Nature, 561(7721), 57–62. doi:10.1038/s41586-018-0459-6

Van Dijk, K. R., Hedden, T., Venkataraman, A., Evans, K. C., Lazar, S. W., & Buckner, R. L. (2010). Intrinsic functional connectivity as a tool for human connectomics: theory, properties, and optimization. Journal of Neurophysiology, 103(1), 297–321. doi:10.1152/jn.00783.2009

van Strien, N. M., Cappaert, N. L. M., & Witter, M. P. (2009). The anatomy of memory: an interactive overview of the parahippocampal-hippocampal network. Nature Reviews Neuroscience, 10(4), 272–282. doi:10.1038/nrn2614

Witter, M. P., & Amaral, D. G. (1991). Entorhinal cortex of the monkey: V. Projections to the dentate gyrus, hippocampus, and subicular complex. Journal of Comparative Neurology, 307(3), 437–459. doi:10.1002/cne.903070308

Witter, M. P., & Amaral, D. G. (2021). The entorhinal cortex of the monkey: VI. Organization of projections from the hippocampus, subiculum, presubiculum, and parasubiculum. Journal of Comparative Neurology, 529(4), 828–852. doi:10.1002/cne.24983

Witter, M. P., Doan, T. P., Jacobsen, B., Nilssen, E. S., & Ohara, S. (2017). Architecture of the Entorhinal Cortex A Review of Entorhinal Anatomy in Rodents with Some Comparative Notes. Frontiers in Systems Neuroscience, 11(46). doi:10.3389/fnsys.2017.00046

Wyss, J. M., & Van Groen, T. (1992). Connections between the retrosplenial cortex and the hippocampal formation in the rat: a review. Hippocampus, 2(1), 1–11. doi:10.1002/hipo.450020102

Zeineh, M. M., Holdsworth, S., Skare, S., Atlas, S. W., & Bammer, R. (2012). Ultra-high resolution diffusion tensor imaging of the microscopic pathways of the medial temporal lobe. Neuroimage, 62(3), 2065–2082. doi:10.1016/j.neuroimage.2012.05.065

